# Genetic Basis of Alternative Polyadenylation is an Emerging Molecular Phenotype for Human Traits and Diseases

**DOI:** 10.1101/570176

**Authors:** Lei Li, Yipeng Gao, Fanglue Peng, Eric J. Wagner, Wei Li

## Abstract

Genome-wide association studies have identified thousands of non-coding variants that are statistically associated with human traits and diseases. However, functional interpretation of these variants remains a major challenge. Here, we describe the first atlas of human <u>3’</u>-UTR alternative polyadenylation (APA) <u>Q</u>uantitative <u>T</u>rait <u>L</u>oci (3’QTLs), i.e. ∼0.4 million genetic variants associated with APA of target genes across 46 Genotype-Tissue Expression (GTEx) tissues from 467 individuals. APA occurs in approximately 70% of human genes and substantively impacts cellular proliferation, differentiation and tumorigenesis. Mechanistically, 3’QTLs could alter polyA motifs and RNA-binding protein binding sites, leading to thousands of APA changes. Importantly, 3’QTLs can be used to interpret ∼16.1% of trait-associated variants and are largely distinct from other QTLs such as eQTLs. The genetic basis of APA (3’QTLs) thus represent a novel molecular phenotype to explain a large fraction of non-coding variants and to provide new insights into complex traits and disease etiologies.

**Highlights:**

- The first atlas of human 3’QTLs: ∼0.4 million genetic variants associated with alternative polyadenylation of target genes across 46 tissues from 467 individuals
- 3’QTLs could alter polyA motifs and RNA-binding protein binding sites
- 3’QTLs can be used to interpret ∼16.1% of trait-associated variants
- Many disease-associated 3’QTLs contribute to phenotype independent of gene expression

## INTRODUCTION

Genome-wide association studies (GWAS) have identified thousands of genetic variants associated with quantitative traits and common diseases. However, the vast majority of variants occur in non-coding regions, thus posing a significant challenge for elucidating the molecular mechanisms by which these variants contribute to diseases and phenotypes. Toward this end, researchers have proposed employing several molecular phenotypes, such as expression quantitative trait loci (eQTLs) (Consortium et al., 2017) (i.e., genetic variants associated with expression of one or more genes), for the functional interpretation of GWAS loci. Although these molecular phenotypes can be informative and in many cases are thought to impact the transcription of nearby genes, the roles of a large fraction of trait-associated non-coding variants remain unexplained (Gamazon et al., 2018).

Alternative polyadenylation (APA) has emerged as a new paradigm of post-transcriptional regulation for most human genes. By employing different polyA sites, genes can either shorten or extend the 3’UTRs that contain *cis*-regulatory elements such as miRNAs or RNA-binding protein (RBP) binding sites (Mayr, 2017). APA can affect the stability and translation efficiency of target mRNAs and the cellular localization of proteins (Tian and Manley, 2016). The diverse landscape of polyadenylation can significantly impact both normal development and development of diseases such as cancer (Mayr, 2018). The importance of polyadenylation was demonstrated in our earlier work (Masamha et al., 2014), which identified *Nudt21* as a key APA regulator that functions as a tumor-suppressor gene in glioblastoma. The results of our more recent study suggested that 3’UTR shortening in breast cancer represses tumor-suppressor genes in *trans* by disrupting competing endogenous (ceRNA) crosstalk (Park et al., 2018).

In addition to their association with gene expression, genetic variations were identified as critical regulatory factors for APA in individual genes in certain cell lines (Yoon et al., 2012). Moreover, APA-associated genetic changes have been linked to the development of multiple disease states, including cancer (Stacey et al., 2011), α-thalassemia (Higgs et al., 1983), facioscapulohumeral muscular dystrophy (van der Maarel et al., 2011), bone fragility (Fahiminiya et al., 2015), neonatal diabetes (Garin et al., 2010), and systemic lupus erythematosus (SLE) (Hellquist et al., 2007) (Graham et al., 2007a). For example, a single SNP (rs10954213) within the 3’UTR of *IRF5* can alter 3’UTR length and affect mRNA stability (Graham et al., 2007), which can further contribute to susceptibility to SLE. Aside from these few examples, the roles of genetic determinants of APA in various human tissues and their association with phenotypic traits and diseases have not been systematically examined.

To obtain insights into the genetic basis of APA regulation in human tissues, we used the DaPars2 (Dynamic analyses of alternative polyadenylation from RNA-seq) algorithm to construct an atlas of tissue-specific human APA events using 7, 655 RNA-seq data coupled with whole-genome sequencing genotype data derived from 46 tissues from 467 individuals in the Genotype-Tissue Expression (GTEx) project. In total, we identified 426,433 genetic variants associated with tissue-specific APA (3’QTLs) sufficient to explain approximately 16.1% of human traits and diseases. Collectively, the results of our study indicate that 3’QTLs represent an emerging molecular phenotype for the interpretation of a significant portion of human genetic variants found outside of coding regions.

## RESULTS

### The atlas of human 3’QTLs: ∼0.4 million genetic variants associated with APA of target genes across 46 tissues from 467 individuals

To detect global APA events in primary human tissues, we used our DaPars2 algorithm to retrospectively and directly identify APA events using 7,655 standard RNA-seq data spanning 46 tissue types from the GTEx v7 project. The multi-sample DaPars2 regression framework calculates a Percentage of Distal polyA site Usage Index (PDUI) for each gene in each sample. The PDUI values were then further normalized after correction of known and inferred technical covariates using probabilistic estimation of expression residuals (PEER) (Stegle et al., 2012). We then used Matrix eQTL to identify genetic variations associated with differential 3’UTR usage in each tissue (Shabalin, 2012) (see ‘Methods’ section for details) (Figure S1). Using a false-discovery rate (FDR) threshold of ≤0.05, we identified 426,433 *cis*-3’QTLs distributed across 46 tissues that were associated with 11,697 genes, representing approximately 51% of annotated genes. The tissues with the highest numbers of 3’QTLs tended to have larger sample sizes (Table S1). This strong association with sample size suggests that more APA events (Figure 1A) and 3’QTLs (Figure 1B) will be discovered when additional RNA-seq datasets become available.

**Figure 1.**
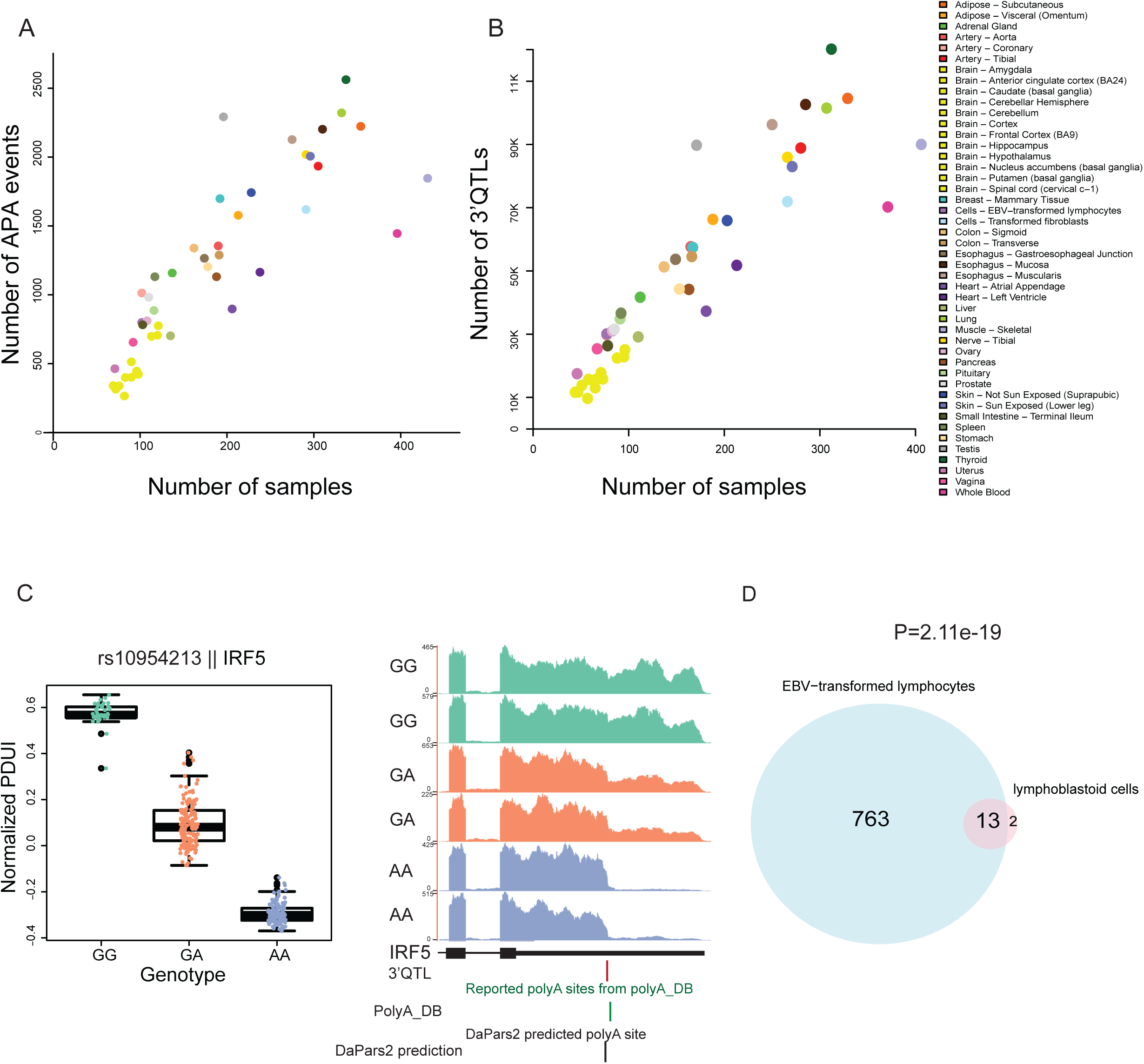
Atlas of genetic variations associated with 3’UTR usage across 46 human tissues. (A) Distribution of the number of APA events with sample size across different tissues. Each color code represents a specific tissue of origin. (B) Distribution of the number of significant 3’QTLs (FDR≤0.05) with tissue sample size. (C) An example 3’QTL (rs10954213) strongly associated with *IRF5* 3’UTR usage in whole blood. Left panel shows the distribution of normalized PDUIs in each genotype. Each dot in the box plot represents the normalized PDUI value of a particular sample. Right panel shows the RNA-seq coverage track for the *IRF5* 3’UTR. The bottom four tracks show RefSeq gene structure, 3’QTL location, reported polyA site location, and DaPars2 prediction. (D) Venn diagram comparison of predicted APA genes from GTEx EBV-transformed lymphocytes and literature-reported APA genes from lymphoblastoid cell lines.

To evaluate the performance of our 3’QTL detection with the current sample size, we compared the detected 3’QTLs with previously reported SNPs associated with variations in 3’UTR usage. Although previous studies of APA events associated with altered gene expression have been limited to a few cell types, such as lymphoblastoid cells, our approach recaptured many of these ‘experimentally validated’ 3’QTLs. For example, the strong association (FDR: 7.26e-159) between SNP rs10954213 and the alternative 3’UTR of *IRF5* (Graham et al., 2007b), a transcription factor involved in multiple immune process, could be detected in our 3’QTL analysis of whole-blood RNA-seq data (Figure 1C). Interestingly, we also found that this genetic effect on *IRF5* was shared with 22 other tissues, suggesting that our multi-tissue context analysis of this locus could aid further investigations of how *IRF5* variants contribute to autoimmune diseases (Figure S2). Of the 17 previously reported SNPs associated with 15 APA genes identified within lymphoblastoid cell lines, our 3’QTL analysis recaptured 13 of the 15 (87%) APA genes in Epstein-Barr virus (EBV)-transformed lymphocytes (Figure 1D). This observation demonstrates that the currently available datasets are useful for capturing most 3’QTLs in human tissues. These results also suggest that the ∼0.4 million 3’QTLs we identified provide an extensive view of how genetic variants are associated with 3’UTR usage across multiple human tissues and expands the number of 3’QTLs by several orders of magnitude compared to all previously reported APA-associated SNPs.

### Genetic variants are associated with APA of HLA-class genes

To investigate the global distribution of 3’QTLs across the human genome, we used Manhattan plots to visualize the locations of 3’QTLs with their associated *P*-values (Figure 2A). Extensive and significant 3’QTL were distributed across each chromosome. Importantly, previously reported APA genes were readily detected, including *IRF5* (Graham et al., 2007b), *ERAP1* (Zhernakova et al., 2013), *THEM4* (Zhernakova et al., 2013), *EIF2A* (Yoon et al., 2012), and *DIP2B* (Yoon et al., 2012); however, the majority of the 3’QTL genes represented novel events. Among these novel 3’QTL genes, several are particularly noteworthy, including *CHURC1*, which encodes a zinc-finger transcriptional activator important in neuronal development (Sheng et al., 2003), and *TPSAB1*, which encodes α-tryptase and reportedly plays a role in a multisystem disorder such as irritable bowel syndrome caused by elevated basal serum tryptase levels (Lyons et al., 2016) (Figure 2B).

**Figure 2.**
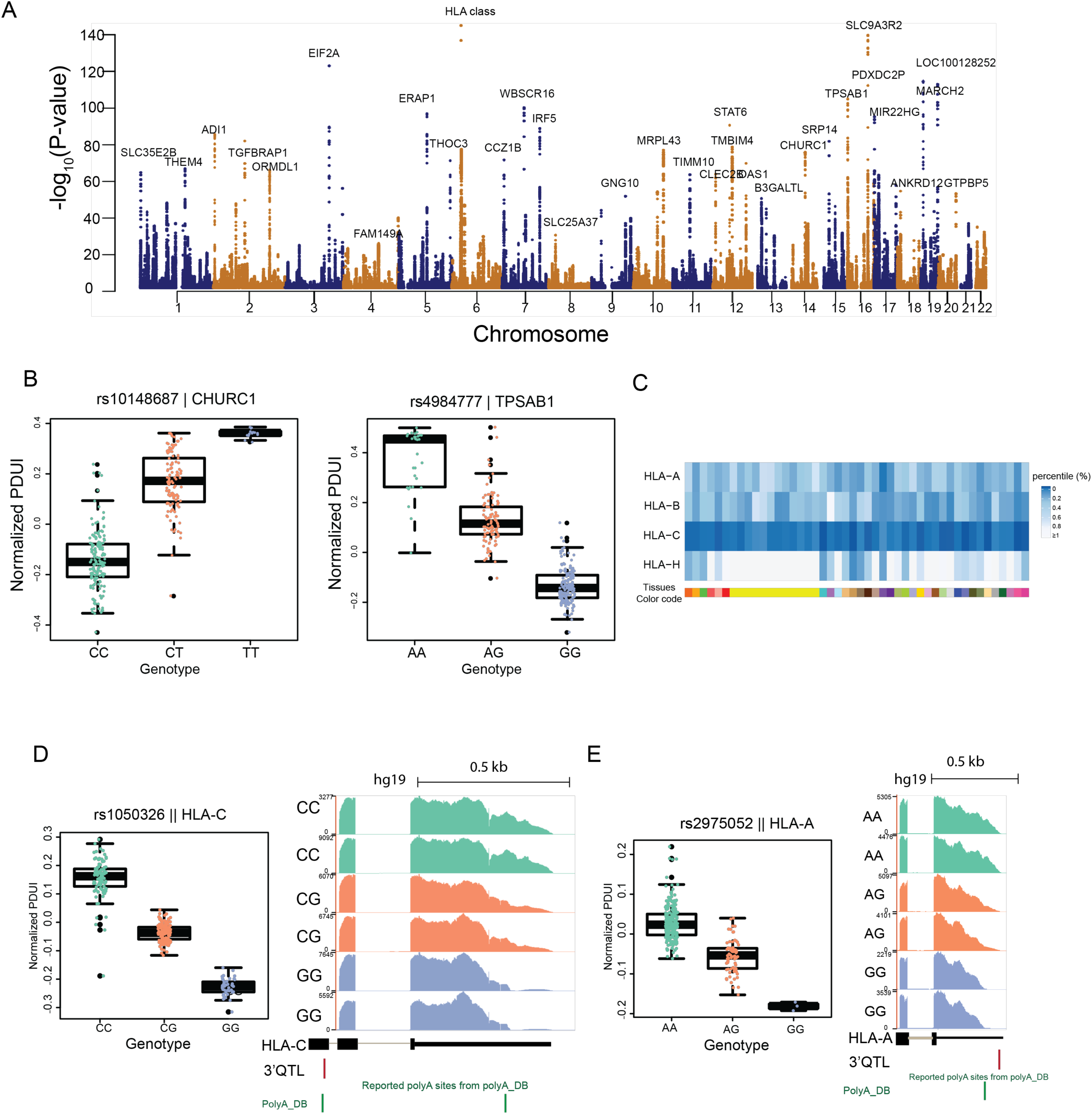
Genetic variations affect 3’UTR usage in human tissues. (A) Manhattan plot of 3’QTL in tissue adipose subcutaneous. The X-axis represents chromosomes locations, and the y-axis shows the −log_10_(*P*-value). Representative APA genes are highlighted. (B) Examples of significant 3’QTL-associated genes in tissue adipose subcutaneous. The plot shows the distribution of normalized PDUIs in each genotype for the genes *MARCH2* and *THOC3*. Each dot represents a normalized PDUI value from a particular sample. (C) Heatmap showing the percentile of *P*-value rank for each HLA-class I gene across human tissues. (D) Example 3’QTL (rs1050326) associated with 3’UTR usage in *HLA-C*. (E) Example 3’QTL (rs2975052) associated with 3’UTR usage in *HLA-A*.

Interestingly, we identified a ‘hotspot’ for 3’QTLs on chromosome 6, and this hotspot was strongly associated with the APA of a human leukocyte antigen (HLA)-class gene. Proteins encoded by HLA gene clusters play essential roles in antigen presentation, and variations in these clusters have been linked to autoimmune diseases (Dendrou et al., 2018). Although genetic variants correlated with HLA gene expression have been extensively studied (Dendrou et al., 2018), there are no published reports describing the influence of *cis*-acting regulatory variants on 3’UTR usage in HLA-class I genes. We conducted a global examination of HLA-class I genes and found extensive genetically associated HLA 3’UTR changes across multiple human tissues (Figure 2C). For example, a variation near the intronic polyA site of *HLA-C* was strongly associated with differential 3’UTR usage (Figure 2D). In the class I HLA gene *HLA-A*, the adenine within the 3’UTR was found to be mutated to guanine, leading to usage of a shortened 3’UTR (Figure 2E). These findings indicate that genetic variants exist that impact 3’UTR APA events within HLA genes and underscore how such variants can impact other genes in a manner distinct from simply inducing a change in expression.

### Complete map of tissue-specific APA usage and 3’QTLs

Prior work indicated that APA events are often regulated in a tissue-specific manner (Zhang et al., 2005) (Weng et al., 2016) (Lianoglou et al., 2013). Although compelling, these previous studies examined only a few tissue types with a relatively small sample size. Here, we employed our computational approach to identify significant APA events across 46 tissues. Overall, we found striking differences in polyA site usage across human tissues (Table S3). We observed that each of 13 distinct brain substructures tends to harbor longer 3’UTR isoforms, whereas several other tissues, such as EBV-transformed lymphocytes, whole blood, and testis, tend to express shorter 3’UTRs (Figure 3A). These results are in agreement with previous findings from analyses of 3’-seq datasets derived from five different tissues (Lianoglou et al., 2013). However, a major strength of our results is that they represent an 8-fold increase in the number of tissues analyzed, thus providing the most complete map of tissue-specific APA usage to date.

**Figure 3.**
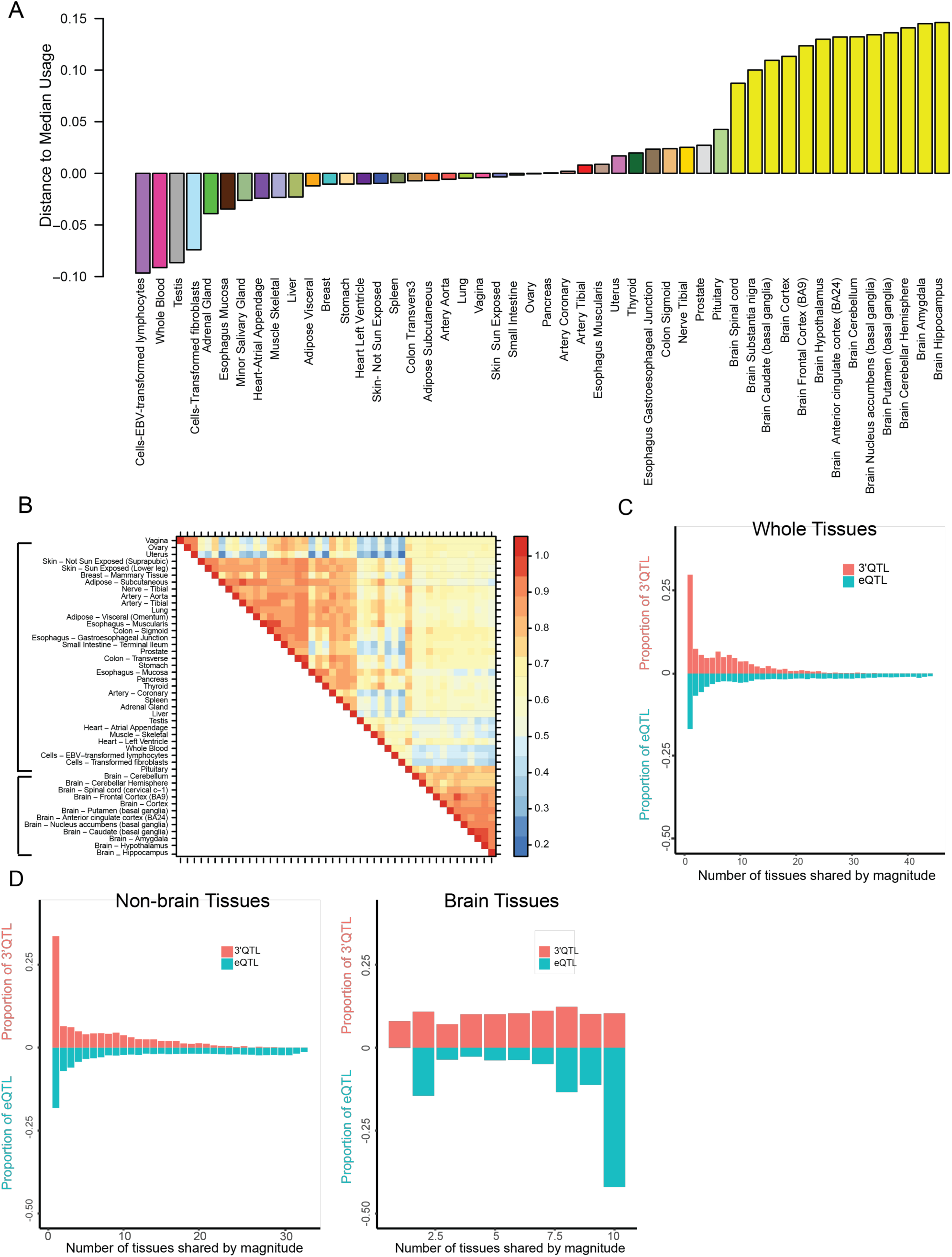
Tissue-specific APA usage and 3’QTL. (A) Distribution of sorted tissue overall 3’UTR scores. Each color code indicates a tissue of origin. The X-axis shows tissue names, and the y-axis indicates the distance of the 3’UTR score for each tissue from the median value across all tissues. (B) Pairwise sharing by 3’QTL magnitude among tissues. Lead 3’QTL were considered, and the proportion of 3’QTL shared with the same sign and within the 2-fold effect size in the other tissue was calculated. Brackets show brain and non-brain tissues. (C) Histograms showing the proportion of mashr-estimated tissues in which the lead 3’QTL/eQTL are shared by magnitude among 46 tissues, considering all tissues, as well as (D) non-brain and brain tissues. Sharing by magnitude means the QTL have the same sign and similar effect size (within 2-fold).

To examine how *cis*-regulatory elements contribute to APA events in a tissue-specific or tissue-sharing manner, we used multivariate adaptive shrinkage (mash) to estimate the effect size of 3’QTLs shared across 46 tissues (Urbut et al., 2018). Correlation values for each tissue were calculated based on the effect sign (effect in the same direction) and magnitude (effect in the same direction and within a 2-fold effect-size change) of 3’QTLs. This analysis revealed that human tissues cluster into two major groups in an unbiased fashion: brain tissues and non-brain tissues (Figure 3B). In addition, we also noted that some biologically related tissues grouped together within “non-brain” tissues, such as the uterus/vagina/ovary group and the colon/stomach group (Figure S3). The pattern of 3’QTLs sharing suggests the possibility of developmental and functional similarities between different tissues. In addition, we found that although 85.9% of tissues had 3’QTLs with the same sign, only 15.7% were shared with 3’QTLs of a similar magnitude. Compared with eQTL shared among tissues (85% of tissues sharing by sign and 36% of tissues sharing by magnitude) (Urbut et al., 2018), 3’QTL exhibited similar sign effects (Figure S4) but a much lower degree of magnitude effects (Figure 3C and 3D). Considered collectively, these observations suggest that 3’QTLs exhibit greater tissue specificity than eQTLs.

### 3’QTLs are distinct from other molecular QTLs

To characterize the relationships between different QTLs, we classified 3’QTLs and eQTLs across 46 tissue types according to the functional category defined in SnpEff (Cingolani et al., 2012). As expected, we found that 3’QTLs were significantly enriched in 3’UTRs (P=1.74e-18) or located within 5 kb downstream of genes (P=7.8e-05), whereas eQTLs were significantly enriched within gene promoters/upstream regions (P=1.45e-26) or with 5’UTRs (P=2.66e-24) (Figure 4A). This observation is consistent with the metagene analysis encompassing the relative position distribution of 3’QTL or eQTL over their associated genes (Figure 4B). Furthermore, 3’QTL differ markedly from splicing-QTL, which are enriched primarily within gene bodies (Li et al., 2016). Interestingly, the structures of 3’QTL-associated genes and eQTL-associated genes differed considerably. Compared with eQTL-associated genes, 3’QTL-associated genes harbored comparable 5’UTRs but much longer CDS (P = 6.71e-25) and 3’UTR lengths (P = 6.21e-105) (Figure 4C). Considered collectively, the results of these analyses suggest that 3’QTL represent a novel type of molecular QTL that differs substantially from other previously defined QTLs.

**Figure 4.**
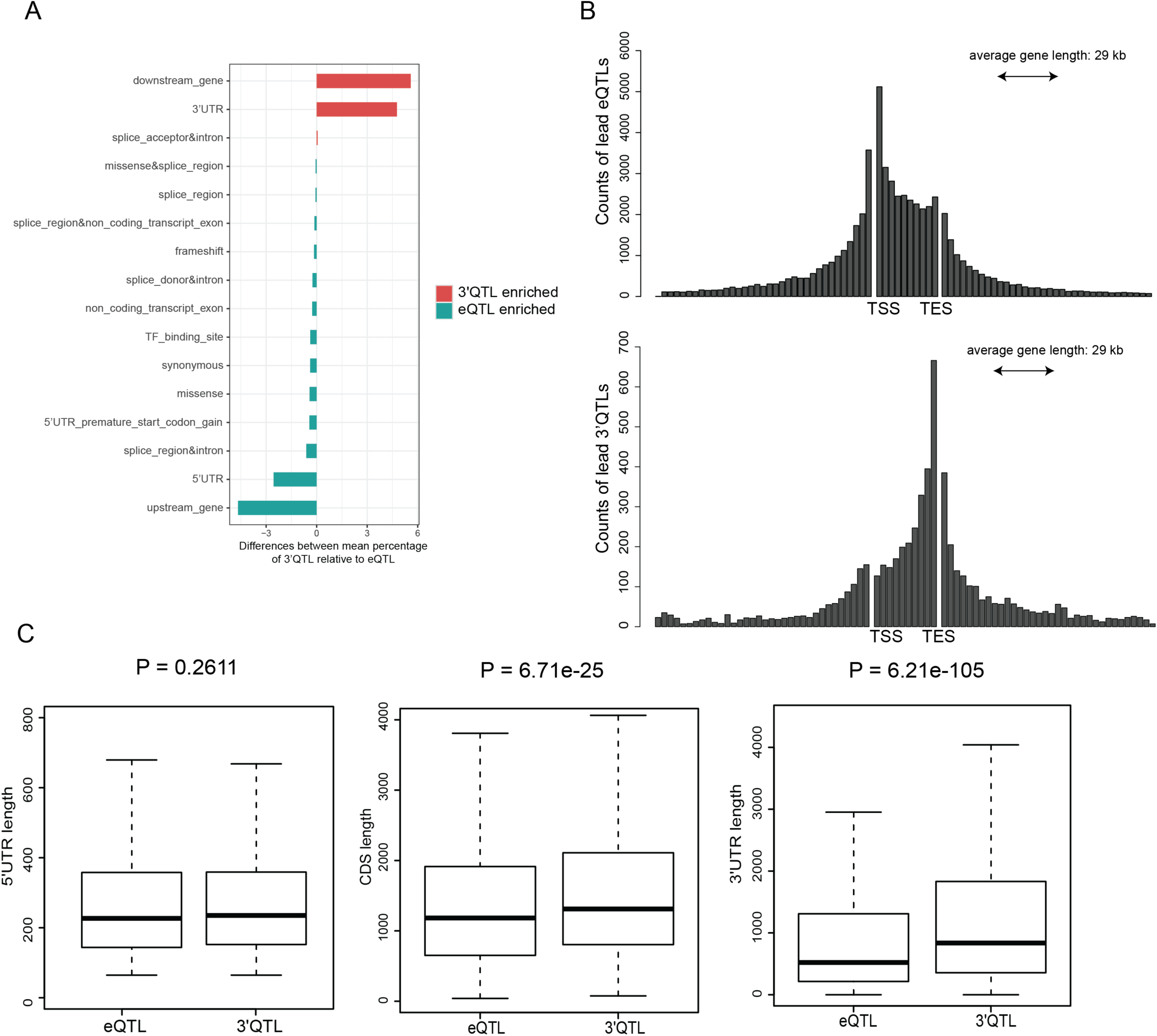
3’QTL represent a new type of molecular QTL. (A) Differences between mean percentage of 3’QTLs and eQTLs among different annotations. Color indicates the statistical significance of the difference, with red indicating 3’QTL-enriched annotations with FDR≤0.01 and green indicating eQTL-enriched annotations with FDR≤0.01. (B) Relative distance between eQTLs and 3’QTLs and their associated APA genes. TSS represent transcription start site; TES represent transcription end site. (C) Genomic length was compared for 3’QTL-associated genes and eQTL-associated genes. *P*-values were calculated using the Student’s *t* test.

### Alteration of the polyA motif is associated with APA

Next, we investigated the potential mechanisms through which genetic variations contribute to APA events. We hypothesized that some 3’QTLs alter the motifs important for 3’-end processing of transcripts. Alteration of polyadenylation motifs can produce distinct mRNA isoforms with 3’UTRs of differing length (Yoon et al., 2012) (Thomas and Sætrom, 2012). However, only a few cases have been reported from a limited number of cell lines. To systematically examine the prevalence of polyA motif–altered 3’QTLs in human populations, we extracted significant 3’QTLs (FDR ≤ 0.05) located within 50 bp upstream of annotated polyA sites compiled from the PolyA database (Lee et al., 2007), UCSC, Ensembl, and RefSeq gene annotations. We performed motif searches based on 15 common polyadenylation motif variants. In total, we identified 2,241 3’QTLs with the potential to alter polyA motifs and generate alternative 3’UTR lengths in the associated genes across 46 human tissues (Figure 5A) (Table S4). For example, a change in SNP rs1130319 from the reference A allele to the C allele, which impairs the canonical polyA motif AATAAA, was found to correlate with the preferred usage of the cryptic polyadenylation site in the *ADI1* 3’UTR (Figure 5B). In another case, G to A change in rs3211995 creates a strong polyadenylation signal (AATAAA) from the weak non-canonical GATAA motif in the 3’-end of *SLC9A3R2*, which was found to correlate with a shift to an mRNA isoform with a longer 3’UTR (Figure 5C). Collectively, the results of our analyses suggest that detectable APA events result from the alteration of polyA motifs in thousands of 3’QTLs.

**Figure 5.**
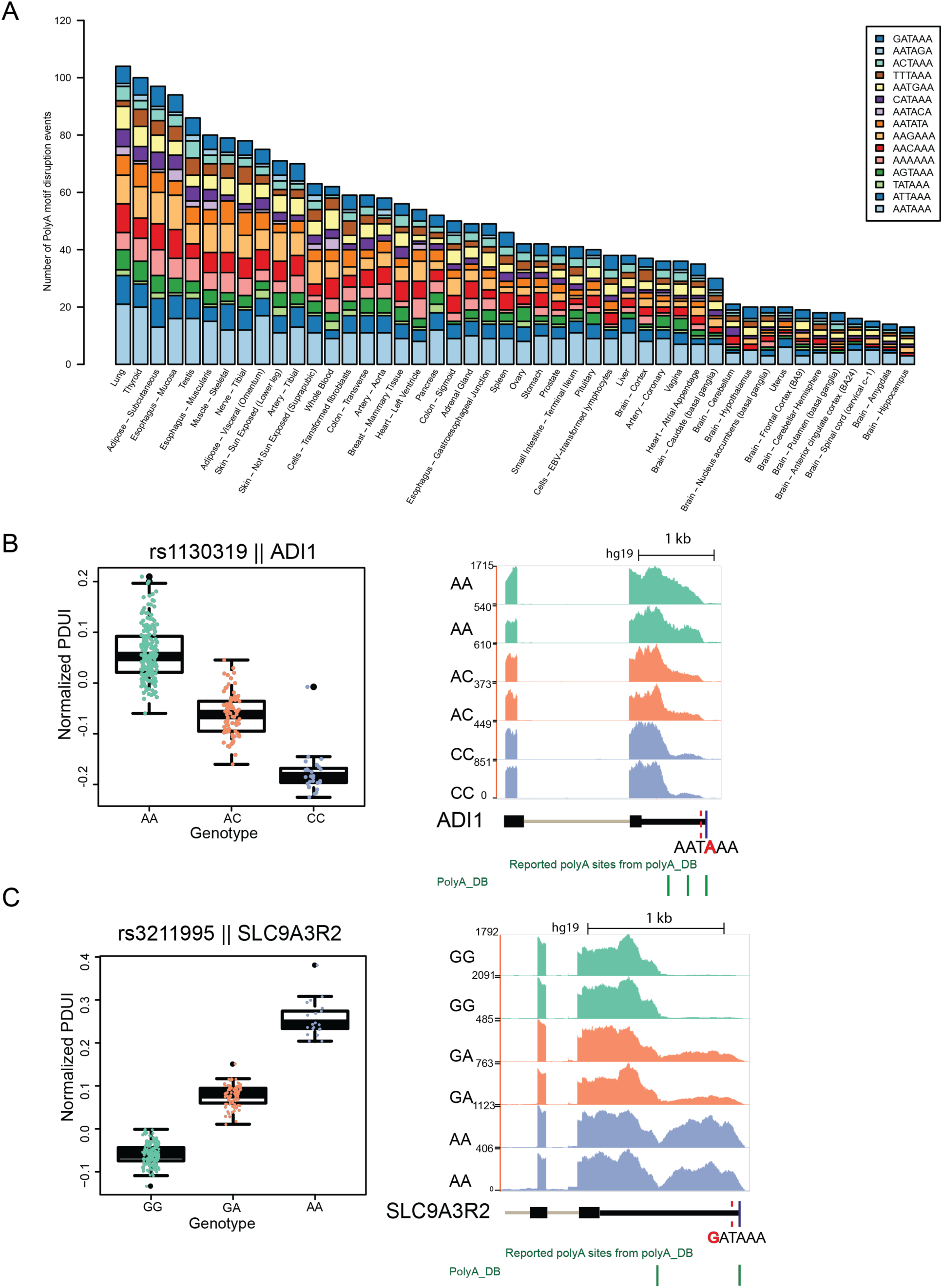
3’QTLs can alter polyA motifs in human tissues. (A) Summary of polyA motifs altered by 3’QTLs across human tissues. The X-axis shows tissue names, and the y-axis lists the number of 3’QTLs that disrupted polyA motifs. (B) Boxplot showing the significant correlation of the 3’QTLs rs1130319 with *ADI1* APA events in each genotype. Each dot represents a normalized PDUI value from a particular sample. Coverage plot illustrates that this SNP could disrupt the polyadenylation motif. The red dotted line in the Ref gene structure indicates the location of 3’QTLs. PolyA motif is shown with the 3’QTL highlighted in red. (C) Boxplot showing that 3’QTL rs3211995 is strongly correlated with *SLC9A3R2* 3’UTR change in each genotype. Coverage plot illustrates that this SNP could “create” a canonical polyadenylation motif.

### Alteration of RBP binding sites is associated with APA

Alteration of polyA motifs can explain only a small percentage of 3’QTLs, suggesting that most 3’QTLs affect APA via other mechanisms. To clarify this issue, we analyzed the extent to which 3’QTLs interfere with either the transcriptional or post-transcriptional regulation of target genes. First, we used DeepBind (Alipanahi et al., 2015) to evaluate the enrichment of 927 binding motifs of 538 DNA-binding proteins and RBPs in 3’QTLs of each tissue and used randomly shuffled 3’QTL as a control. We identified 125 motifs that were significantly enriched in 3’QTLs, 17 of which were common between at least 20% of the tissues examined (Figure 6A). These motif-associated genes included several known polyadenylation factors, such as *PABP* (Matoulkova et al., 2012), *CPEB4* (Bava et al., 2013; Matoulkova et al., 2012), and *HNRNPC*, which was recently described as an APA regulator (Gruber et al., 2016).

**Figure 6.**
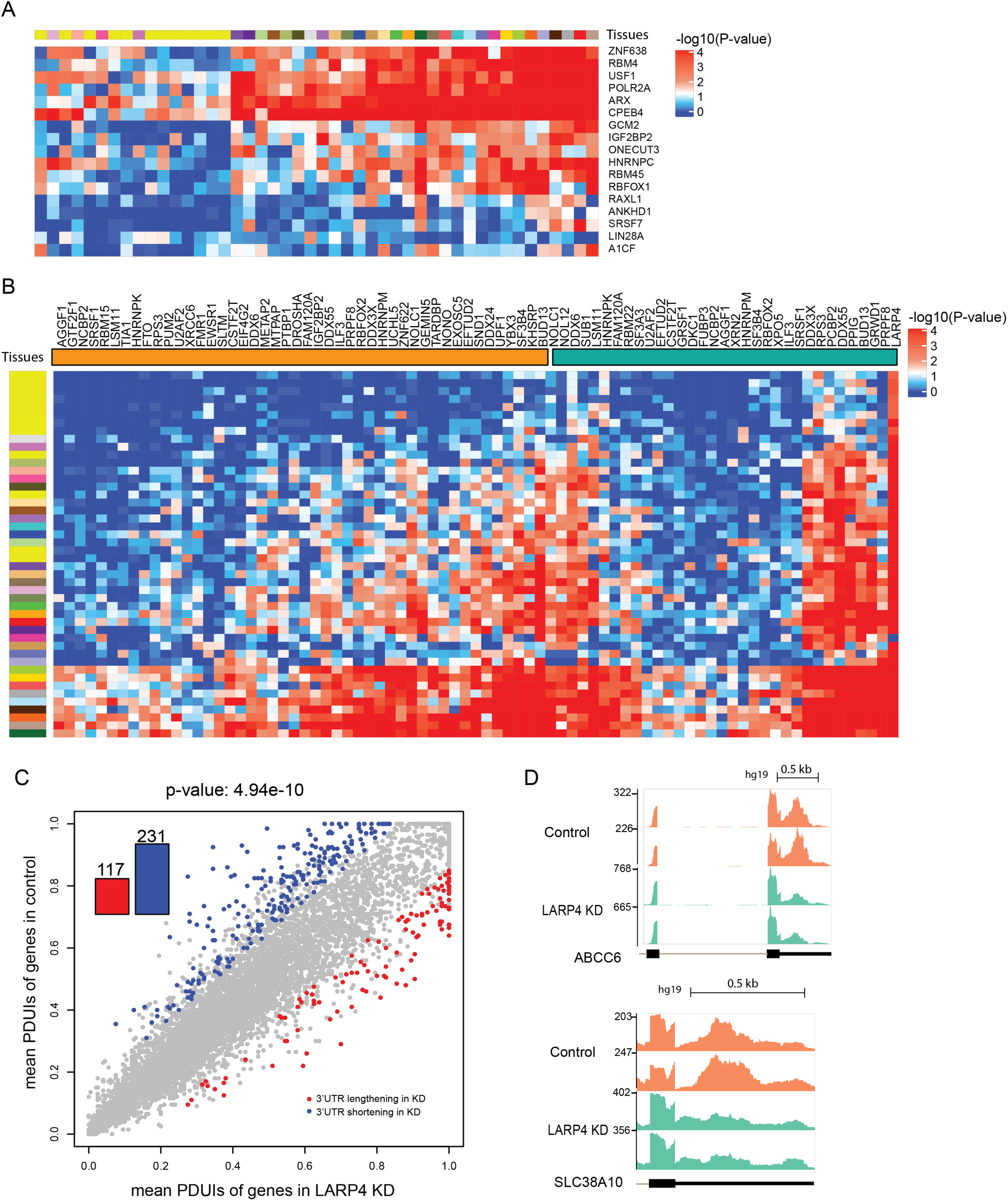
Alteration of binding sites of RNA-binding proteins can affect 3’UTR length. (A) Heatmap of 3’QTL significance for RNA motifs from DeepBind in each tissue. Values in the heatmap indicate enrichment of 3’QTL in RNA motifs compared to randomly shuffled SNPs. Top bar shows the color code for each tissue. (B) Heatmap of 3’QTL significance for RNA-binding proteins from ENCODE in each tissue. Left bar shows the color code for each tissue, and top color bar represents K562 and HepG2 cell lines, separately. Values in the heatmap represent the degree of enrichment of 3’QTLs in RNA-binding protein binding peaks compared to the control. (C) Scatterplot of PDUIs in shRNA knockdown of *LARP4* and the control. The shift toward proximal sites upon shRNA knockdown was significant (*P*=4.94e-10, binomial test). (D) Examples of genes in which the 3’UTR was altered upon *LARP4* knockdown.

We then analyzed RBP CLIP-seq data available from the ENCODE project (Dominguez et al., 2018). This dataset is particularly useful given that 81.2% of RBPs are not included in the DeepBind resource. We examined whether 3’QTLs are significantly enriched within the CLIP binding peaks of each RBP in comparison with a random sequence dataset. We identified 79 RBPs that preferentially bind to regions containing 3’QTLs, and notably, these included several polyA factors, such as *CSTF*, in addition to many splicing factors (Figure 6B). Consistent with a potential functional significance, these splicing factors were previously been linked to alternative 3’UTR usage (Bava et al., 2013) (Muller-McNicoll et al., 2016).

We found that one tumor-suppressor gene, *LARP4* (La ribonucleoprotein domain family member 4) selectively bound to 3’QTL-containing regions across most tissues. LARP4 is an RBP that binds to the polyA tail of mRNA molecules (Yang et al., 2011) and regulates mRNA translation; however, no role in APA regulation has been reported. Our observation that *LARP4* binding involves regions enriched with 3’QTLs suggests that *LARP4* may be a novel APA regulator. To test this hypothesis, we re-analyzed *LARP4* shRNA-knockdown gene expression datasets from the ENCODE project (Dominguez et al., 2018). Differential 3’UTR usage was calculated using DaPars (Xia et al., 2014). Interestingly, we found that depletion of *LARP4* led to significant 3’UTR shortening (P=4.94e-10), suggesting that *LARP4* is a novel APA regulator (Figure 6C and 6D). Considered collectively, our results suggest that alteration of RBP binding sites could explain how 3’QTLs contribute to APA events.

### A significant proportion of disease heritability can be explained by 3’QTLs

The GWAS approach is commonly used to associate genetic variants with complex human traits and diseases. However, it is often difficult to explain how these genetic variations, particularly non-coding variations, contribute to specific phenotypes. We thus hypothesized that 3’QTL represent a novel molecular phenotype that could be used to interpret non-coding variants, particularly those located around 3’UTRs. Here, we compiled GWAS summary statistics regarding 23 common human disease and traits from published studies (Table S5) and evaluated the enrichment of lead 3’QTLs within trait-association GWAS SNPs in each tissue using fgwas (Pickrell, 2014). We found there is an enrichment of 3’QTLs within 54.1% tissue-trait pairs (Figure S5). To further compare with lead eQTLs enrichment for these traits, we observed that, overall, eQTLs had a larger effect than 3’QTLs in the tissue-trait pairs examined (49.3%). However, in several case, we also found that 3’QTLs exhibited stronger enrichment of GWAS SNPs than eQTLs (Table S6), for example with Alzheimer disease, rheumatoid arthritis, and type 2 diabetes. Notably, many of the 3’QTLs enriched for trait associations were from tissues relevant to their respective diseased state, such as the spleen and thyroid for type 2 diabetes, the brain cortex for Alzheimer disease, and the pituitary for rheumatoid arthritis (Figure 7A).

**Figure 7.**
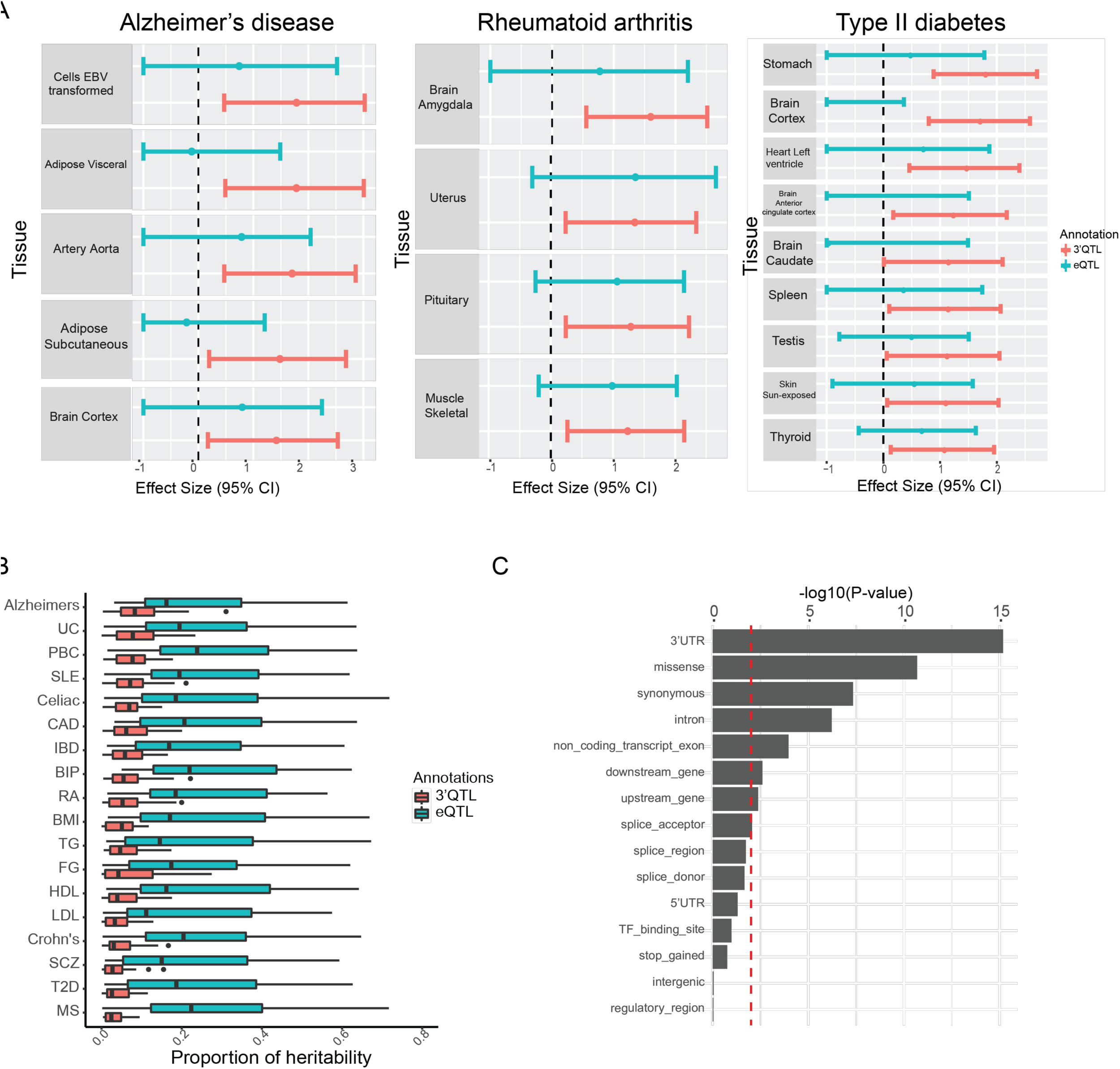
Association of 3’QTLs with human GWAS diseases. (A) Enrichment in tissues with a greater 3’QTL enrichment compared to eQTL for Alzheimer disease, rheumatoid arthritis, and type 2 diabetes. The effect size represents the QTL enrichment result calculated based on fgwas results. (B) Partitioned heritability plot of the percentage of SNP heritability explained (x-axis) for 28 traits (y-axis) by eQTLs and 3’QTLs. (C) Functional term enrichment of 3’QTL-associated GWAS SNPs relative to non-3’QTL–associated GWAS SNPs. Red line indicates a *P*-value of 0.01.

To further quantify the proportion of regulatory variations in heritability for each trait, we conducted a partitioned heritability analysis using trait summary statistics– based LDSR (linkage disequilibrium score regression) (Finucane et al., 2015). Of the traits examined, the median range of SNP heritability explained by 3’QTL and eQTL was 2–9% and 11–24% per trait, respectively. Notably, 3’QTL were particularly effective in explaining a large proportion of heritability for several autoimmune diseases, such as ulcerative colitis, primary biliary cholangitis, and Alzheimer disease. For some diseases, such as multiple sclerosis, 3’QTLs contributed little to heritability (Figure 7B).

As GWAS summary statistics were limited to a few common diseases and traits, we broadened our analysis to identify the most strongly associated 3’QTLs among 34,087 GWAS SNPs for 2,183 traits derived from the NHGRI GWAS catalog (MacArthur et al., 2017). In total, 2,751 (8%) GWAS SNPs were either common to or in strong linkage disequilibrium (LD) (r^2^ ≥ 0.8) with lead 3’QTLs. To better characterize 3’QTL-associated GWAS SNPs, we analyzed the enrichment of functional terms as defined in the GWAS catalog of 3’QTL-associated GWAS SNPs compared with non-3’QTL– associated GWAS SNPs. GWAS SNPs in LD with 3’QTLs were more enriched with 3’UTR (P=5.55e-16), missense, synonymous, and intron variants (Figure 7C). These results suggest that 3’QTLs explain a significant proportion of disease-associated variants.

### Many disease–co-localizing 3’QTL contribute to phenotypes independent of gene expression

The strong LD of genetic variants with disease-associated loci does not necessarily imply a causal relationship. Some eQTLs are known causal variants in human disease and traits (Musunuru et al., 2010). We therefore investigated to what extent 3’QTL may function as causal variants for human phenotypes using co-localization analysis (Giambartolomei et al., 2014) of 15 complex diseases and traits with known minor allele frequencies in order to identify 3’QTL sharing the same putative causal variants with trait-associated signals. In total, 801 trait-associated variants co-localized with either eQTLs or 3’QTLs signals. Consistent with previous results (Consortium et al., 2017), 57% of the trait-associated variants co-localized with eQTL in one or more tissues. Interestingly, 16.1% of trait-associated variants co-localize with 3’QTLs in at least one tissue (Figure 8A). To determine whether 3’QTL co-localizing genes are also shared with eQTL, we compared 1,267 3’QTL-trait gene pairs with eQTL-trait gene pairs across all tissues and found that 85% (1,075/1,267) of the 3’QTL–co-localized genes were specific for 3’QTL but not eQTL (Figure 8B) (Table S7).

**Figure 8.**
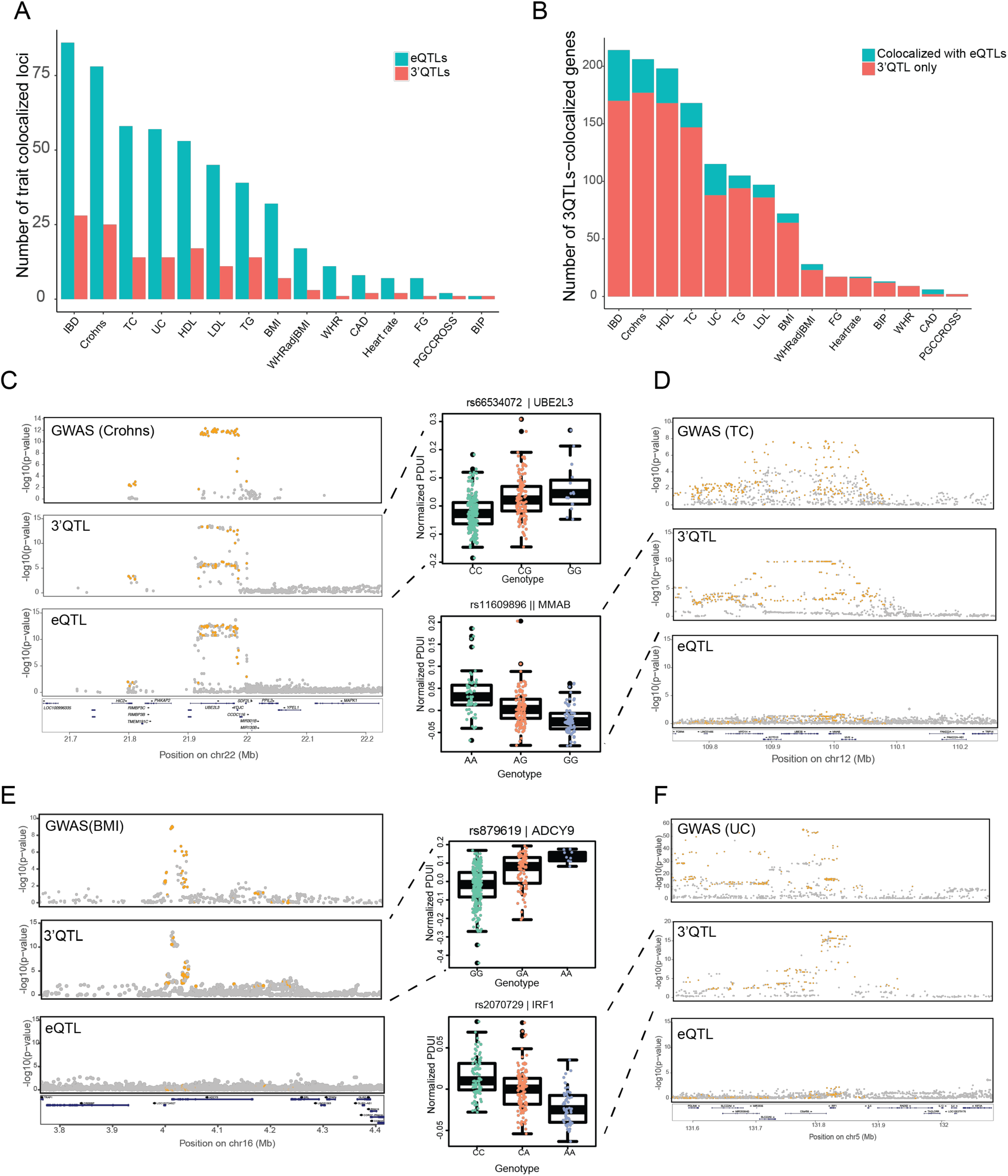
Co-localization of 3’QTL with complex trait–associated loci. (A) Total number of co-localized GWAS signals (y-axis) for each of 15 traits (x-axis) across human tissues. (B) The number of 3’QTL co-localizing genes, colored based on whether the gene also co-localized with eQTLs. (C) Co-localization map of Crohn disease with 3’QTLs in whole blood. Boxplot shows that 3’QTLs are strongly associated with 3’UTR usage in *UBE2L3*. Orange indicates 3’QTLs shared with GWAS SNPs. (D) Co-localization map of total cholesterol levels trait with 3’QTLs and eQTLs in liver tissue. (E) Co-localization map of body mass index trait with 3’QTLs and eQTLs in skeletal muscle tissue. (F) Co-localization map of ulcerative colitis with 3’QTLs and eQTLs in transformed fibroblasts.

*UBE2L3* is a representative example of the 15% of genes that were 3’QTL– /eQTL–co-localizing. *UBE2L3* is an E2 ubiquitin-conjugating enzyme that promotes activation of NF-κB signaling in immune responses (Lewis et al., 2015). The rs66534072 locus in *UBE2L3* is associated with the gene’s expression level and confers risk for SLE (Wang et al., 2012). However, the mechanism regulating how these genetic variants affect gene expression remains unknown. We determined that *UBE2L3* is subject to APA and can exhibit dynamic 3’UTR usage in different individuals. Moreover, the lead 3’QTL SNP rs66534072 was significantly correlated with 3’UTR usage in *UBE2L3* (Figure 8C). Specifically, the C allele was associated with *UBE2L3* 3’UTR mRNA shortening, whereas the G allele was associated with 3’UTR lengthening. We examined the tissues in which rs66534072 serves as a 3’QTL for *UBE2L3* and found that most are autoimmune disease related (Figure S6).

Most 3’QTL-trait co-localized gene pairs are specific to the 3’QTLs itself but not to eQTLs. For instance, 288 3’QTLs associated with *MMAB* 3’UTR usage were directly correlated with total cholesterol level GWAS loci on chromosome 12 (Figure 8D). Similarly, variants on chromosome 16 associated with body mass index also co-localized with 3’QTLs that regulate 3’UTR length change in *ADCY9* (Figure 8E). We also observed a strong co-localization pattern between 3’QTLs of *IRF1* and the significant GWAS loci of multiple autoimmune diseases, including ulcerative colitis, Crohn disease, and inflammatory bowel disease (Figure 8F). *IRF1* is induced by IFN-γ signaling and promotes innate and acquired immune responses (Kano et al., 2008). In contrast, except in muscle-skeletal tissues, there was no strong association between eQTL and *IRF1* expression. Co-localization analyses of muscle-skeletal tissue revealed no co-localization pattern between disease-associated loci and *IRF1* eQTLs. In contrast, co-localization patterns for *IRF1* 3’QTL with autoimmune diseases were identified in multiple tissues, including transformed fibroblasts (PP4=0.97; PP4, posterior probability of a model with one shared causal variant), pancreas (PP4=0.96), aorta (PP4=0.93), esophageal mucosa (PP4=0.90), vagina (PP4=0.82), and coronary artery (PP4=0.80). These results suggest that *IRF1*-associated 3’QTL—more so than eQTL—explain most of the effects of *IRF1* variations on these diseases. Collectively, our data suggest that many 3’QTLs contribute to human diseases and traits independent of regulating gene expression.

## DISCUSSION

We defined 3’QTL as representing a novel human molecular phenotype responsible for alternative 3’UTR usage associated with genetic variations. By re-analyzing large-scale GTEx data using our DaPars2 algorithm, we identified 11,697 APA genes and ∼0.4 million 3’QTLs across 46 human tissues. In contrast to other molecular QTLs, such as eQTLs, 3’QTLs are highly enriched within 3’UTRs. Mechanistically, 3’QTL likely induce changes in 3’UTR usage by either modulating the strength of polyadenylation signal motifs or RBP binding sites. eQTLs are known to be important molecular traits associated with human phenotypic variations. We demonstrated that 3’QTL represent a novel molecular trait that, unexpectedly, is of similar importance as eQTL in contributing to phenotypic variations in human populations.

We found that 3’QTLs explain a large proportion of trait heritability. Co-localization analyses found that 16.1% of trait-associated loci co-localize with one or more 3’QTLs in human tissues. Furthermore, very few of the 3’QTL–co-localizing trait-associated loci overlapped with eQTLs, indicating that 3’QTLs and eQTLs are largely mutually independent. We speculate that eQTL-independent 3’QTLs regulate the stability, translation, or cellular localization of target genes, independent of regulating gene expression. Collectively, the results of our in-depth analyses of the genetic influence of APA events in 46 human tissues are illustrative of genomic “dark matter” beyond coding regions and thus suggest novel interpretations of how natural variations shape human phenotypic diversity and tissue-specific diseases.

## AUTHOR CONTRIBUTIONS

L.L. and W.L. conceived and supervised the project. L.L., and Y. G. performed the data analysis. L.L., F.P., E.J.W. and W.L. interpreted the data and wrote the manuscript.

## COMPETEING INTERESTS

The authors declare no competing financial interests.

## ACKNOWLEDGEMENTS

We thank Lei Hou, Zhe Cui and members of the Li lab for helpful discussions. This work was supported by the US National Institutes of Health (R01HG007538, R01CA193466, R01CA228140).

## SUPPLEMENTARY FIGURE LEGENDS

**Figure S1.**
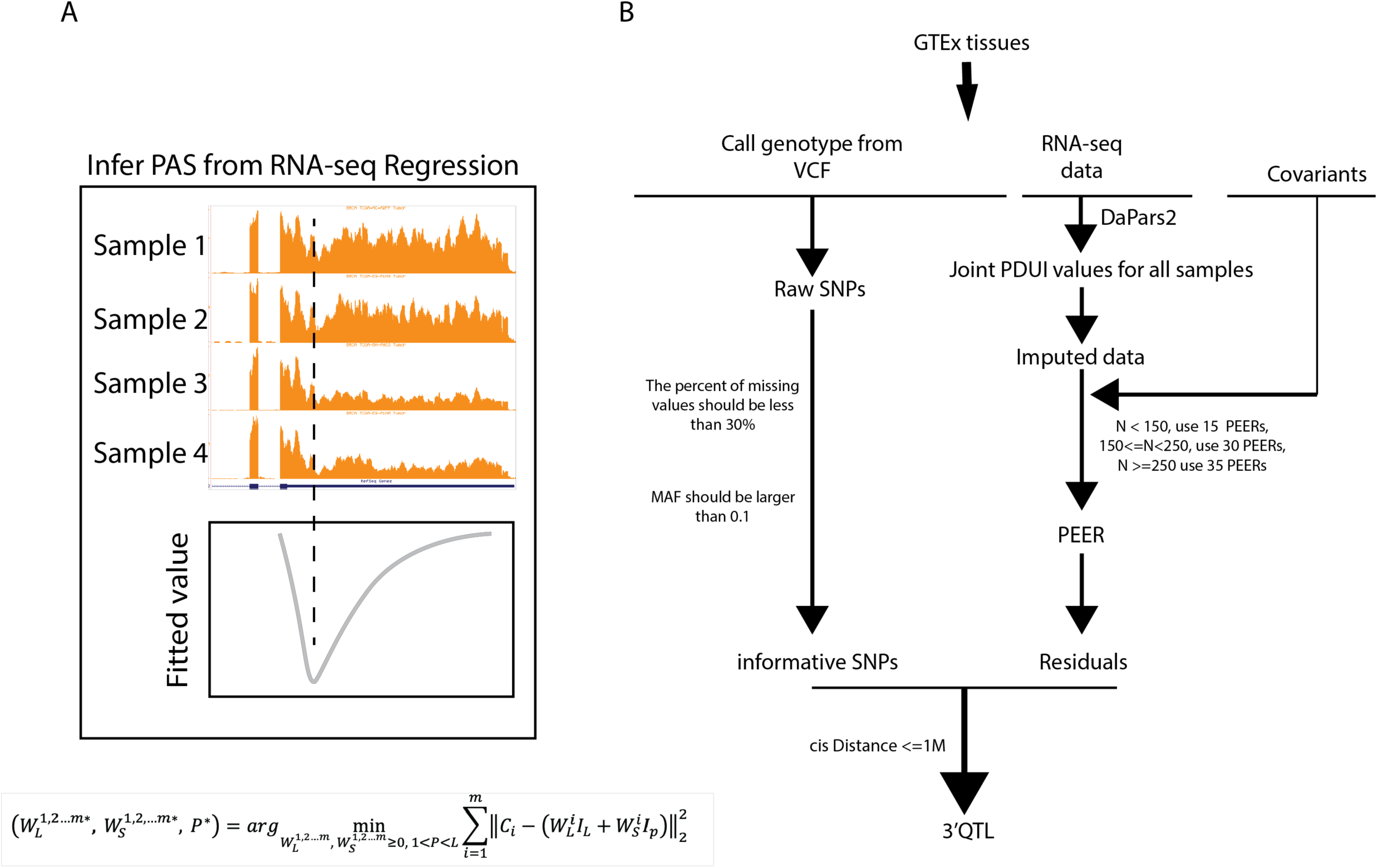
Computational methods to identify 3’QTLs from GTEx data. (A) Statistical framework for identifying APA genes using joint PDUI values from multi-sample RNA-seq data (DaPars2). RNA-seq reads located within 3’UTRs were analyzed together, and a linear regression model was applied to infer the location of *de novo* proximal polyA sites. For each transcript in samples, a PDUI score is assigned. (B) Computational workflow for identifying 3’QTLs from human GTEx data.

**Figure S2.**
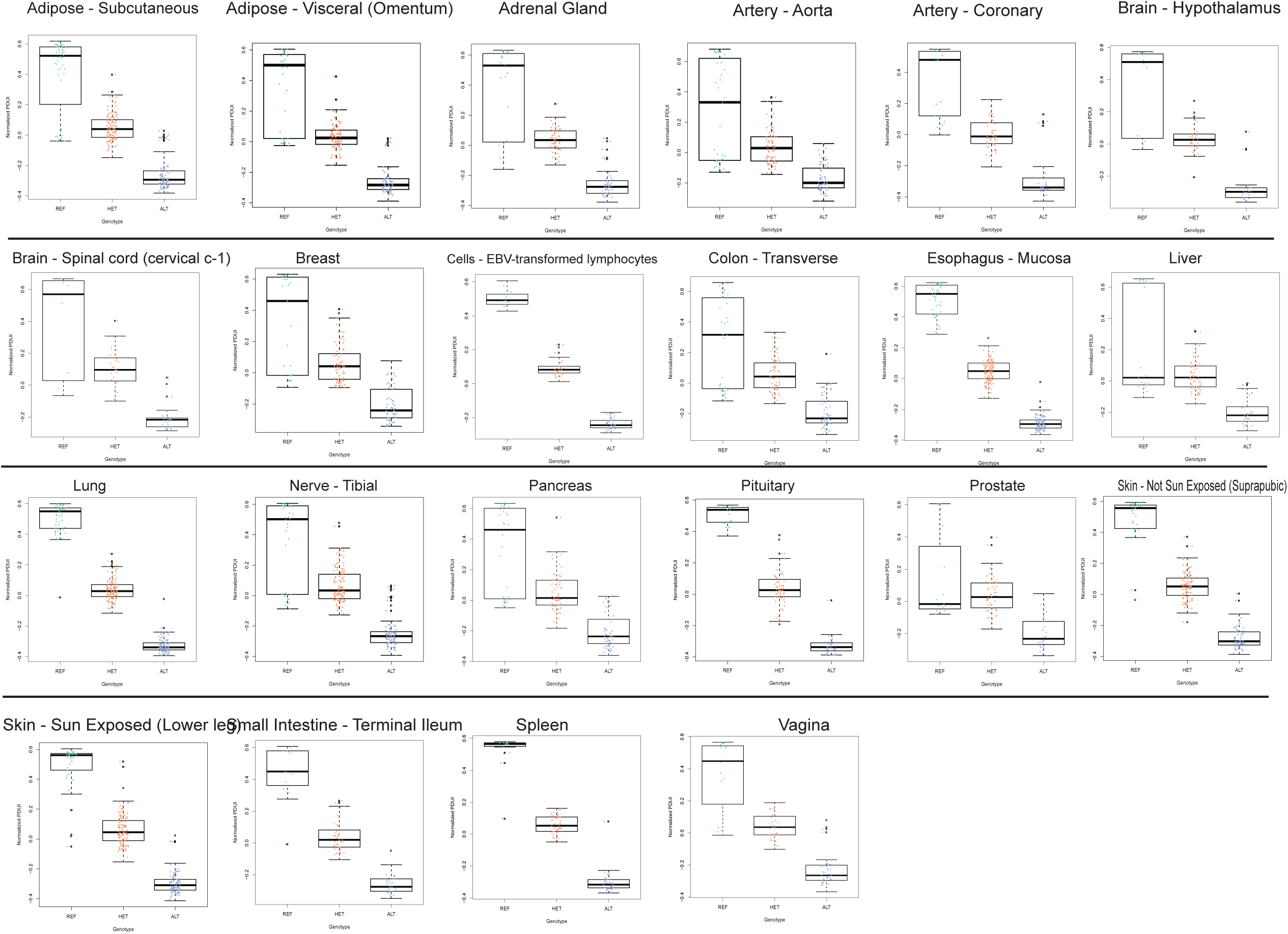
3’QTL rs10954213 is correlated with *IRF5* 3’UTR usage in 22 other human tissues. Each boxplot represents the PDUI difference of *IRF5* in relation to the genotype of the index SNP rs10954213 in each tissue. The X-axis shows the significance of the association, and the y-axis indicates the tissue name. “REF” denotes genotype GG; “HET” denotes genotype GA; and “ALT” denotes genotype AA.

**Figure S3.**
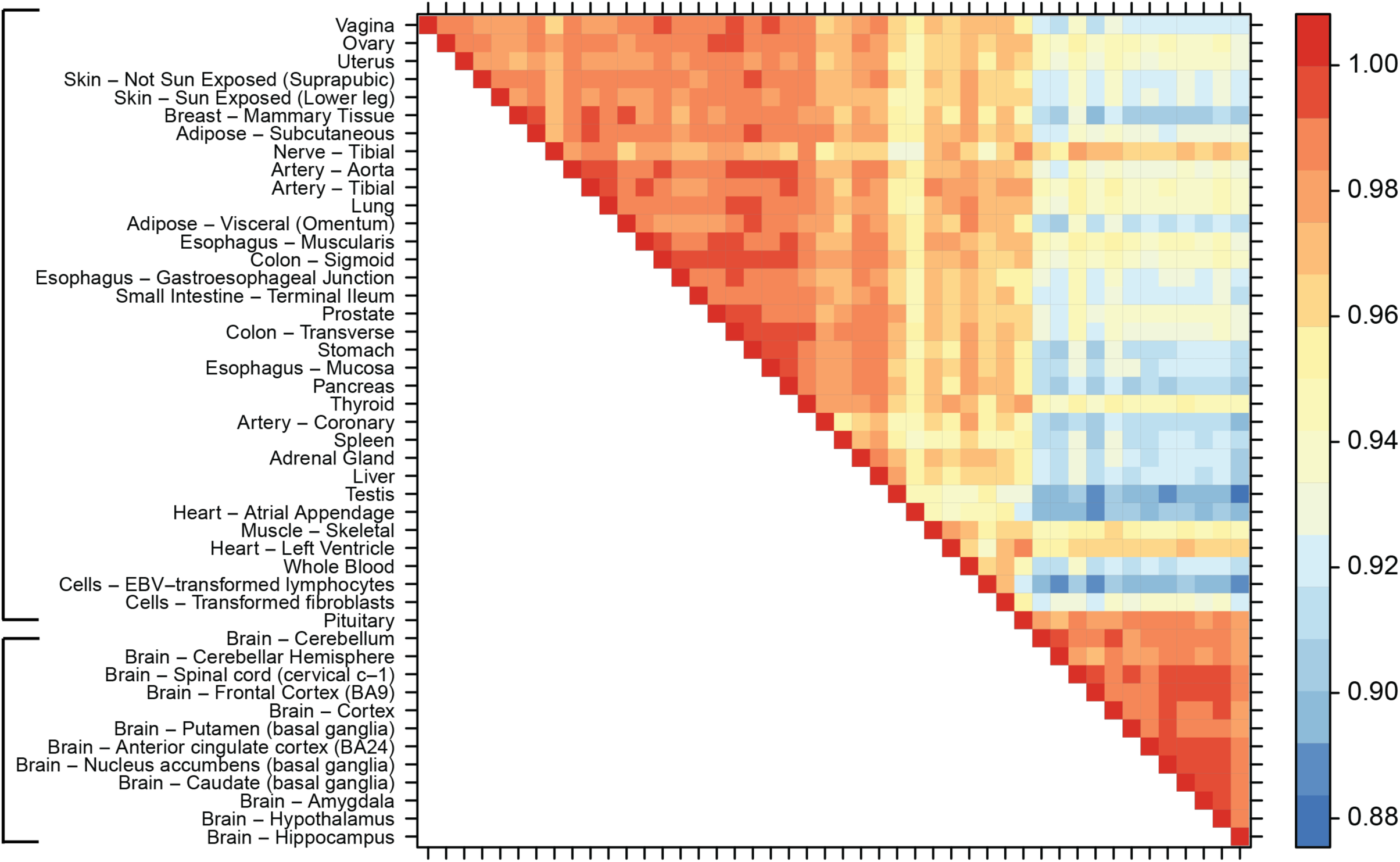
Pairwise sharing by 3’QTL sign among tissues. For each pair of tissues, the proportion of lead 3’QTLs shared in the same direction in the other tissue was calculated. Brackets show brain and non-brain tissues.

**Figure S4.**
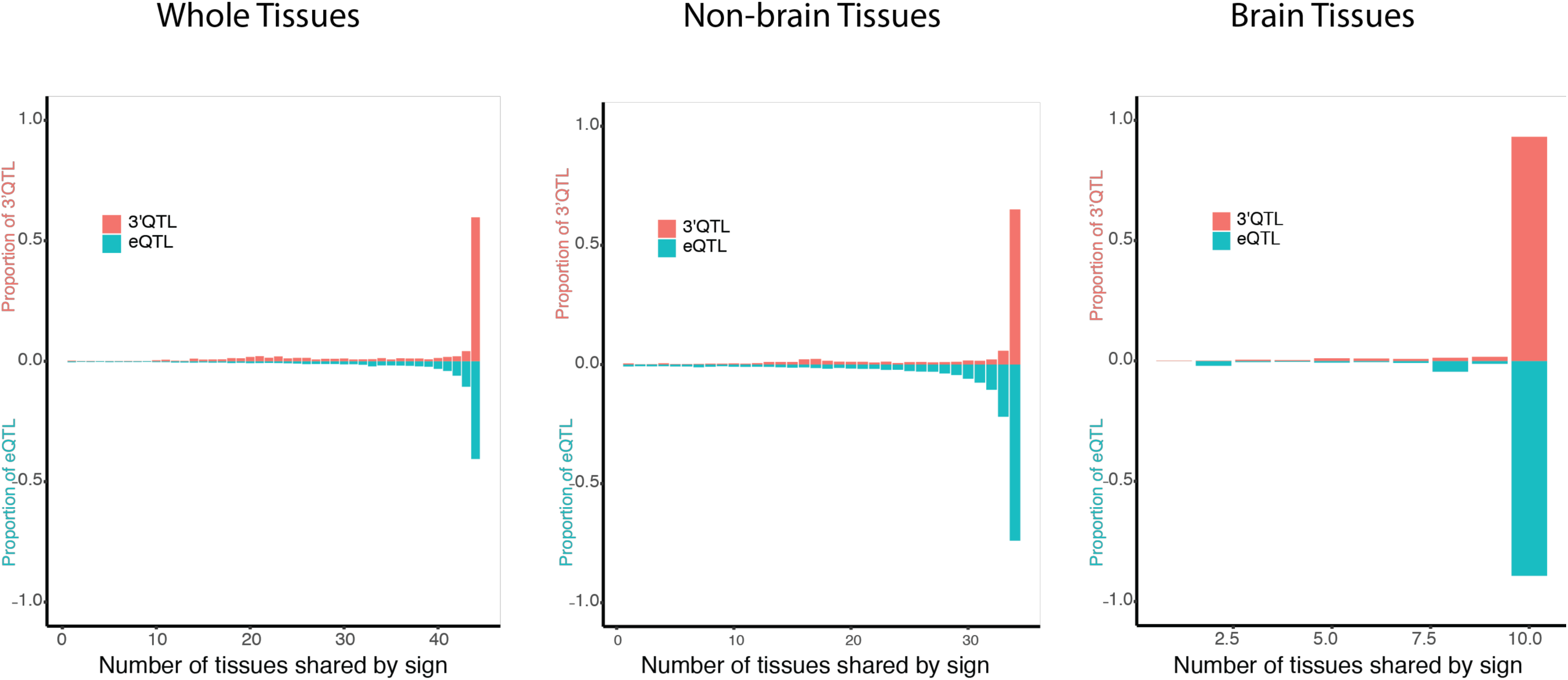
Comparison of pairwise sharing by 3’QTL/eQTL signs. Histograms show the proportion of mash-estimated tissues in which the lead 3’QTL/eQTL are shared by sign among 46 tissues, considering all tissues, as well as non-brain and brain tissues. Sharing by sign means the 3’QTL have the same direction in the mash-estimated effect.

**Figure S5.**
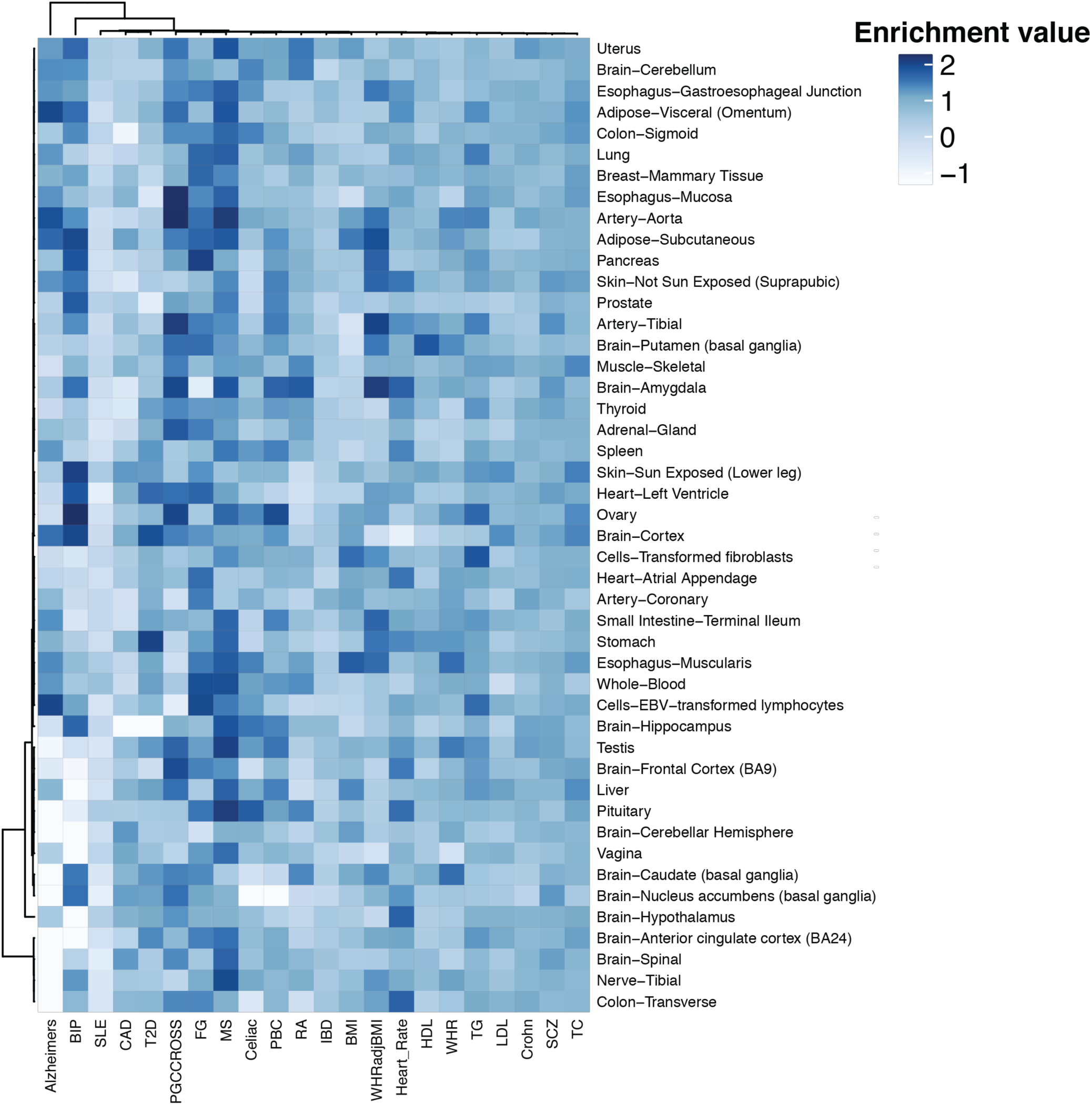
Heatmap of enrichment value for 22 complex traits among 46 tissues using fgwas. The enrichment values (effect size) are calculated using fgwas which quantify the relationship between trait-associated variants and causal 3’QTLs and colored from white to blue. Traits (column) were hierarchical clustered using “median” method. Enrichment values and 95% confidence intervals can be found in Supplementary Table 4.

**Figure S6.**
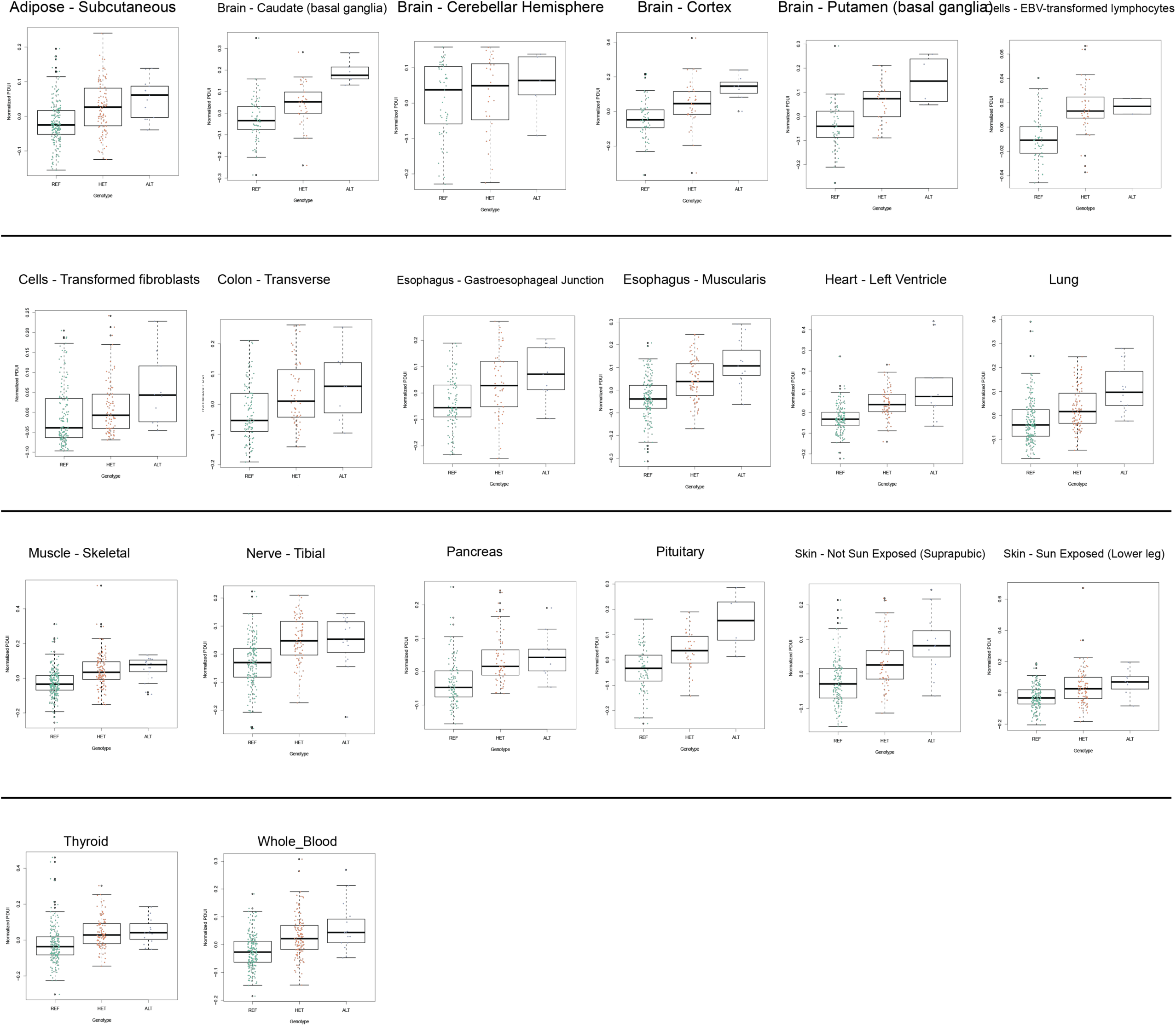
3’QTLs are associated with *UBE2L3* 3’UTR usage in human tissues. Each boxplot represents the correlation of 3’QTL (rs66534072) with *UBE2L3* APA level in each tissue. Each dot in the boxplot represents a PDUI value for *UBE2L3* from a particular sample. “REF” denotes genotype CC; “HET” denotes genotype CG; and “ALT” denotes genotype GG.

## SUPPLEMENTARY TABLE LEGENDS

**Table S1. List of APA and 3’QTLs identified from GTEx data**

APA genes and 3’QTLs identified in this study. Columns include: 1) Tissue Name, 2) Pixel, 3) Colors, 4) Number of 3’QTLs with FDR ≤ 0.05 and 5) Number of APA genes with q-value ≤ 0.05.

**Table S2. List of 3’QTL identified as disrupting polyA motifs**

Table listing each polyA motif disrupted by 3’QTL.

**Table S3. List of human diseases and complex traits examined in this study**

Table listing the different complex human traits and diseases examined, with download URL provided.

**Table S4. fgwas enrichment results**

Table listing the effect size of functional annotations (eQTLs and 3’QTLs) in GWAS signals. Lower_CI and upper_CI represent the lower and upper bound of the confidence interval of the effect size.

**Table S5. List of co-localizing human trait GWAS signals and 3’QTLs**

Table listing genes in which the associated 3’QTL co-localized with GWAS signals. PP0 indicates the null model of no association. PP1 and PP2 indicate the probability that causal variants are either associated with disease signals only or 3’QTL only. PP3 indicates the probability that the genetic effects of disease signals and 3’QTL are independent, and PP4 indicates the probability that disease signals and 3’QTL share causal SNPs.

## Detailed Methods Li et al. 2019

### GTEx data

Raw GTEx RNA-seq and genotype files were obtained from dbGaP release under accession number phs000424.v7.p2. This release consists of 9,777 RNA-seq samples across 54 tissues from 550 individuals. We removed disease tissue cells-leukemia cell line. In addition, seven other tissues, including the cervix endocervix, cervix ectocervix, fallopian tube, bladder, kidney cortex, minor salivary gland, and brain substantia nigra were removed because of small sample size. The related sample description files are available from GTEx Portal (www.gtexportal.org). Individuals not included in the GTEx analyze freeze were also filtered.

### Mapping of GTEx RNA-seq data

Original RNA-seq reads were aligned to the human genome (hg19/GRCh37) using STAR, version 2.5.2b (Dobin et al., 2013), with the following alignment parameters: -- outSAMtype BAM SortedByCoordinate; --outSAMstrandField intronMotif; -- outFilterMultimapNmax 10; --outFilterMultimapScoreRange 1; --alignSJDBoverhangMin 1; --sjdbScore 2; --alignIntronMin 20; and --alignSJoverhangMin 8. The resulting sorted BAM files were converted into bedgraph format using bedtools version 2.17.0 (Quinlan and Hall, 2010).

### DaPars2 analyses

DaPars2 allows expanded DaPars (Xia et al., 2014) analysis of pairwise tumor/normal comparisons to multiple RNA-seq joint analyses, and the dynamics of APA genes can be determined based on a two-normal mixture model.

*Joint Regression model in DaPars2. W*_*L*_ and *W*_*s*_ represent the mean abundance of RefSeq transcripts with distal and proximal polyA sites for sample *i*, respectively, *C*_*i*_ represents the read coverage of sample *i* at single-nucleotide resolution normalized by total sequencing depth; *L* represents the length of the longest 3′ UTR, *p* represents the length of alternative proximal 3′ UTR to be estimated, *I*_*L*_ and *I*_*p*_ represent indicator functions. The expression levels of two transcripts with distal and proximal polyA sites in normal individual samples can be estimated by optimizing this linear regression model. The optimal proximal polyA site *P*\* is selected if it exhibits the minimum deviation between observed and expected read density.

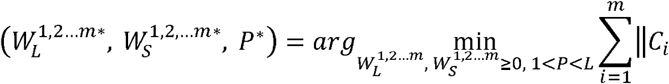

We further calculated the percentage of distal polyA site usage index (PDUI) for each transcript in each sample, with PDUI for each sample defined as:

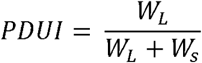

We required that the average normalized reads for each RefSeq 3’ UTR region be >30; otherwise, the PDUI for this transcript would be denoted as a missing value “NA”. For a given tissue, transcripts with missing values in >50% of individuals were removed.

### Covariate estimation

To account for hidden batch effects and other potential covariates in each tissue, we used PEER (Probabilistic Estimation of Expression Residuals) (Stegle et al., 2012), with gender and genotyping platform as the known covariates to estimate a set of latent covariates for PDUI values in each tissue. The number of PEER factors was optimized based on suggestions from the GTEx consortium (Consortium et al., 2017); for tissue sample sizes <150, a PEER factors of 15 was chosen. Thirty PEER factors were chosen if the sample size ranged from 150 to 250, and 35 peer factors were chosen for >250 samples.

### 3’QTL mapping for each tissue

A whole-genome sequencing variants file for 635 individuals was obtained from the GTEx dbGaP phs000424.v7.p2 website under the name ‘GTEx_Analysis_2016-01-15_v7_WholeGenomeSeq_635Ind_PASS_AB02_GQ20_HETX_MISS15_PLINKQC.vcf. gz’, in which 17 samples and all variants that failed to pass the Quality Control step initially defined by the GTEx consortium (Consortium et al., 2017) were removed. The individuals with no RNA-seq data were removed. 3’QTL mapping was performed separately for each tissue. Subset VCF data for each tissue were extracted using bcftools (version 1.3), and only biallelic SNPs were considered. VCF files were transformed into a SNP matrix file with genotyping information included using bioalcidae (Lindenbaum and Redon, 2018). SNPs with a minor allele frequency of <0.01 were filtered. We then tested associations for SNPs within an interval of 1 Mb of the 3’UTR region with normalized PDUI values in each tissue using Matrix eQTL (Shabalin, 2012) in a linear regression framework.

A permutation analysis was conducted to identify significant 3’QTL-associated gene pairs. Individual labels were randomly sampled 1,000 times, and the minimum *P*-value for each SNP and gene after each 3’QTL mapping was recorded. These empirical *P*-values were adjusted using qvalue r package (Storey and Tibshirani, 2003). Genes with a qvalue of <0.05 were considered significant APA genes. All APA gene– associated 3’QTL were subsequently identified with FDR control at 5%.

### Functional annotation of 3’QTL

GTEx eQTL results were obtained from the GTEx portal (www.gtexportal.org). Lead 3’QTL and eQTL SNPs were functionally annotated using SnpEff (Cingolani et al., 2012) version 4.3. Lead QTL were defined as QTL with the minimum *P*-value associated with each APA gene. Each type of annotation was required to have an average of at least 3 assigned QTL for each tissue. The Student’s *t*-test was used to evaluate annotation differences between 3’QTL and eQTL among all tissues. The *P*-values were further adjusted using the Bonferroni correction.

### Tissue 3’QTL sharing and specificity analyses

3’QTL tissue sharing and specificity were analyzed using multivariate adaptive shrinkage (MASH) (Urbut et al., 2019). Briefly, we converted 3’QTL association statistics to mash format. Lead 3’QTL and random SNP sets for each APA gene were extracted from each tissue to calculate MASH priors. A total of 4,470 genes with no missing data in any tissue were retained for training the MASH model. Prior covariance matrices were calculated by FLASH, and the multivariate 3’QTL model was constructed using mashr. Posterior values were computed by applying the trained model to the lead 3’QTL sets.

### Tissue-specific APA score

For each tissue, the mean PDUI value of each transcript was calculated based on the average for all individuals. We subsequently applied a two-component normal mixture model to identify APA genes across multiple tissues. After the regression analysis of the PDUI histogram, the differential PDUI usage index (*DI*), defined as the effective distance normalized by the variance between the two components, was used to quantify PDUI heterogeneity for APA detection:

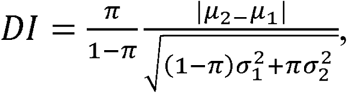

where π represents the proportion of tissues in first group. *µ*_1_ and *µ*_2_: means of the PDUI level of two different modes. The terms *σ*_1_ and *σ*_2_ represent the standard deviations. APA genes were defined as *DI* ≥ 1 and |*µ*_2_ *µ*_1_| ≥ 0.1. The mean value of the identified APA genes in each tissue was considered the overall 3’UTR score for the tissue.

### Meta-gene analysis

We identified the relative location of both eQTL and 3’QTL within 100 kb of the gene by assigning the QTL to distance bins inside or outside of the genes. For bins inside the gene, we set 1/10 of the annotated gene length as the bin size; therefore, each gene was divided into 10 equal-size distance bins. For bins outside the genes, we used an average gene length of ∼29 kb as the bin size and extended the number of bins until it reached the 100-kb cutoff. We then assigned each eQTL and 3’QTL into corresponding distance bins based on chromosome, position, and strand information.

### Identification of polyA motif–disrupting 3’QTL

Significant 3’QTL (FDR ≤ 0.05) located within the 3’UTR of the associated APA gene were intersected with 50 bp upstream of the annotated polyA site compiled from a collection of UCSC Ensemble gene models and PolyADB (Lee et al., 2007). Sequences located 5 nucleotides upstream and downstream of polyA-overlapping 3’QTL were extracted to identify 3’QTL that disrupt polyA motifs.

### Identification of RNA motif–disrupting 3’QTL

We selected significant 3’QTL (FDR ≤ 0.05) located inside the gene region, defined as 3 kb downstream of the transcription start site to the end of the gene. For all 3’QTL, 3 bp of nearby nucleotides were retrieved. We then used DeepBind v0.11 (Alipanahi et al., 2015) to score the 7-mer sequences using 617 pre-built models including 472 transcription factors and 80 RBPs from *Homo sapiens*. For each 7-mer, we kept the top three predictions with a DeepBind score of ≥0.1. To evaluate binding enrichment, we also randomly shuffled the genomic locations of 3’QTL while remaining within the gene region of the genome and on the same chromosome. The Fisher exact test was used to calculate *P*-values for differences between the number of strongly bound 7-mers for 3’QTL and randomly shuffled SNPs.

### Identification of RBP peak–disrupting 3’QTL

CLIP-Seq data were downloaded from the ENCODE data portal (www.encodeproject.org). We restricted our filtering condition to “*Homo sapiens*”, “RNA binding”, and “bed narrowPeak” with genome assembly hg19. Significantly enriched peaks with *P*-values <0.01 were extracted. We retained binding peaks shared by two biological replicates, leaving 166 RBP peak sets from two immortalized cell lines (HepG2 and K562). To examine the associations between the 166 RBP binding sites and 3’QTL, we compared 3’QTL of each tissue with RBP binding peaks using bedtools (Quinlan and Hall, 2010). The statistical significance of resulting overlaps was evaluated using a random permutation approach. We randomly shuffled the genomic locations of 3’QTL while remaining within the gene region and on the same chromosome. Fisher’s exact test was employed to test for enrichment.

### Identification of trait-associated 3’QTL

The RefSNP accessions of 3’QTL were identified using biomaRt package (Durinck et al., 2009). Linkage disequilibrium (LD) datasets for the European population were obtained from Google Genomics (https://console.cloud.google.com/storage/browser/genomics-public-data/linkage-disequilibrium/1000-genomes-phase-3/ldCutoff0.4_window1MB/sub_pop/CEU/?pli=1). GWAS SNPs were obtained from the NHGRI GWAS catalog (MacArthur et al., 2017) (accessed 2018/06/21, v1.0), and 34,087 SNPs were left after removing SNPs with either no dbSNP accession or multiple SNPs associated with one trait. 3’QTL were considered to overlap with disease-associated loci if the lead 3’QTL or its LD tag (r^2^ ≥0.8) mapped with a GWAS SNP.

### FGWAS analyses

GWAS full summary statistics for 23 common traits were compiled from the literature (Table S3). We re-formatted these summary files to contain RefSNP id, chromosomal position, effect size (or OR), standard error, and Z-score. We then applied fgwas (Pickrell, 2014) to these GWAS results to estimate enrichment effects of different genomic annotations within trait-associated GWAS SNPs. Briefly, the genomic regions were annotated in a binary fashion as 3’QTL or eQTL. We used the default parameters (window size=5000, prior variance=0.01, 0.1, 0.5) to run fgwas and adjusted the window size for three other diseases (Crohn disease, Inflammatory bowel disease, celiac disease) to fit the requirements of fgwas.

### Co-localization analyses

We utilized a Bayesian co-localization approach to identify GWAS signals that could exhibit the same genetic effect between eQTL and 3’QTL using coloc R package (Giambartolomei et al., 2014). A total of 15 GWAS full summary statistics were used when the minor allele frequency was available. For each GWAS trait, we extracted the sentinal SNPs, which were defined as GWAS SNPs with a *P*-value of <5×10^-8^ and located at least 1 Mb away from more significant variants. The co-localized signals were searched within a surrounding region of 100 kb of sentinel SNPs. As defined by the coloc method, five posterior probabilities (PPs) were calculated. PP0 represents the null model of no association. PP1 and PP2 represent the probability that causal genetic variants are either associated with disease signals only or 3’QTL only. PP3 represents the probability that the genetic effects of disease signals and 3’QTL are independent, and PP4 represents the probability that disease signals and 3’QTL share causal SNPs. The genes were defined as co-localization events if PP4≥0.75 and PP4/(PP4+PP3) ≤0.9. Region visualization plots were constructed using LocusZoom (Pruim et al., 2010). LD association between reference SNPs and 3’QTL was calculated using PLINK (Purcell et al., 2007).

### Partitioned heritability analysis

The relative contribution of 3’QTL and eQTL to SNP heritability of 28 complex diseases and trait was estimated using ldsc software (Bulik-Sullivan et al., 2015) (https://github.com/bulik/ldsc). We ran LDSR on significant 3’QTL and eQTL variant-gene pairs (FDR≤0.05) in each tissue using default parameters.

## REFERENCES

Alipanahi, B., Delong, A., Weirauch, M.T., and Frey, B.J. (2015). Predicting the sequence specificities of DNA- and RNA-binding proteins by deep learning. Nat Biotechnol 33, 831–838.

Bava, F.A., Eliscovich, C., Ferreira, P.G., Minana, B., Ben-Dov, C., Guigo, R., Valcarcel, J., and Mendez, R. (2013). CPEB1 coordinates alternative 3’-UTR formation with translational regulation. Nature 495, 121–125.

Cingolani, P., Platts, A., Wang le, L., Coon, M., Nguyen, T., Wang, L., Land, S.J., Lu, X., and Ruden, D.M. (2012). A program for annotating and predicting the effects of single nucleotide polymorphisms, SnpEff: SNPs in the genome of Drosophila melanogaster strain w1118; iso-2; iso-3. Fly (Austin) 6, 80–92.

Consortium, G.T., Laboratory, D.A., Coordinating Center-Analysis Working, G., Statistical Methods groups-Analysis Working, G., Enhancing, G.g., Fund, N.I.H.C., Nih/Nci, Nih/Nhgri, Nih/Nimh, Nih/Nida, et al. (2017). Genetic effects on gene expression across human tissues. Nature 550, 204–213.

Dendrou, C.A., Petersen, J., Rossjohn, J., and Fugger, L. (2018). HLA variation and disease. Nat Rev Immunol 18, 325–339.

Dominguez, D., Freese, P., Alexis, M.S., Su, A., Hochman, M., Palden, T., Bazile, C., Lambert, N.J., Van Nostrand, E.L., Pratt, G.A., et al. (2018). Sequence, Structure, and Context Preferences of Human RNA Binding Proteins. Mol Cell 70, 854–867 e859.

Fahiminiya, S., Al-Jallad, H., Majewski, J., Palomo, T., Moffatt, P., Roschger, P., Klaushofer, K., Glorieux, F.H., and Rauch, F. (2015). A polyadenylation site variant causes transcript-specific BMP1 deficiency and frequent fractures in children. Human Molecular Genetics 24, 516–524.

Finucane, H.K., Bulik-Sullivan, B., Gusev, A., Trynka, G., Reshef, Y., Loh, P.R., Anttila, V., Xu, H., Zang, C., Farh, K., et al. (2015). Partitioning heritability by functional annotation using genome-wide association summary statistics. Nat Genet 47, 1228–1235.

Gamazon, E.R., Segre, A.V., van de Bunt, M., Wen, X., Xi, H.S., Hormozdiari, F., Ongen, H., Konkashbaev, A., Derks, E.M., Aguet, F., et al. (2018). Using an atlas of gene regulation across 44 human tissues to inform complex disease- and trait-associated variation. Nat Genet 50, 956– 967.

Garin, I., Edghill, E.L., Akerman, I., Rubio-Cabezas, O., Rica, I., Locke, J.M., Maestro, M.A., Alshaikh, A., Bundak, R., del Castillo, G., et al. (2010). Recessive mutations in the INS gene result in neonatal diabetes through reduced insulin biosynthesis. Proceedings of the National Academy of Sciences 107, 3105–3110.

Giambartolomei, C., Vukcevic, D., Schadt, E.E., Franke, L., Hingorani, A.D., Wallace, C., and Plagnol, V. (2014). Bayesian test for colocalisation between pairs of genetic association studies using summary statistics. PloS Genet 10, e1004383.

Graham, D.S.C., Manku, H., Wagner, S., Reid, J., Timms, K., Gutin, A., Lanchbury, J.S., and Vyse, T.J. (2007a). Association of IRF5 in UK SLE families identifies a variant involved in polyadenylation. Human Molecular Genetics 16, 579-591-591.

Graham, R.R., Kyogoku, C., Sigurdsson, S., Vlasova, I.A., Davies, L.R., Baechler, E.C., Plenge, R.M., Koeuth, T., Ortmann, W.A., Hom, G., et al. (2007b). Three functional variants of IFN regulatory factor 5 (IRF5) define risk and protective haplotypes for human lupus. Proc Natl Acad Sci U S A 104, 6758–6763.

Graham, R.R., Kyogoku, C., Sigurdsson, S., Vlasova, I.A., Davies, L.R.L., Baechler, E.C., Plenge, R.M., Koeuth, T., Ortmann, W.A., Hom, G., et al. (2007c). Three functional variants of IFN regulatory factor 5 (IRF5) define risk and protective haplotypes for human lupus. Proceedings of the National Academy of Sciences of the United States of America 104, 6758–6763.

Gruber, A.J., Schmidt, R., Gruber, A.R., Martin, G., Ghosh, S., Belmadani, M., Keller, W., and Zavolan, M. (2016). A comprehensive analysis of 3’ end sequencing data sets reveals novel polyadenylation signals and the repressive role of heterogeneous ribonucleoprotein C on cleavage and polyadenylation. Genome Res 26, 1145–1159.

Hellquist, A., Zucchelli, M., Kivinen, K., Saarialho-Kere, U., Koskenmies, S., Widen, E., Julkunen, H., Wong, A., Karjalainen-Lindsberg, M.-L., Skoog, T., et al. (2007). The human GIMAP5 gene has a common polyadenylation polymorphism increasing risk to systemic lupus erythematosus. Journal of Medical Genetics 44, 314-321-321.

Higgs, D.R., Goodbourn, S.E.Y., Lamb, J., Clegg, J.B., Weatherall, D.J., and Proudfoot, N.J. (1983). α-Thalassaemia caused by a polyadenylation signal mutation. Nature 306, 306398a306390.

Kano, S., Sato, K., Morishita, Y., Vollstedt, S., Kim, S., Bishop, K., Honda, K., Kubo, M., and Taniguchi, T. (2008). The contribution of transcription factor IRF1 to the interferon-gamma- interleukin 12 signaling axis and TH1 versus TH-17 differentiation of CD4+ T cells. Nat Immunol 9, 34–41.

Lee, J.Y., Yeh, I., Park, J.Y., and Tian, B. (2007). PolyA_DB 2: mRNA polyadenylation sites in vertebrate genes. Nucleic Acids Res 35, D165–168.

Lewis, M.J., Vyse, S., Shields, A.M., Boeltz, S., Gordon, P.A., Spector, T.D., Lehner, P.J., Walczak, H., and Vyse, T.J. (2015). UBE2L3 polymorphism amplifies NF-kappaB activation and promotes plasma cell development, linking linear ubiquitination to multiple autoimmune diseases. Am J Hum Genet 96, 221–234.

Li, Y.I., van de Geijn, B., Raj, A., Knowles, D.A., Petti, A.A., Golan, D., Gilad, Y., and Pritchard, J.K. (2016). RNA splicing is a primary link between genetic variation and disease. Science 352, 600– 604.

Lianoglou, S., Garg, V., Yang, J.L., Leslie, C.S., and Mayr, C. (2013). Ubiquitously transcribed genes use alternative polyadenylation to achieve tissue-specific expression. Genes Dev 27, 2380–2396.

Lyons, J.J., Yu, X., Hughes, J.D., Le, Q.T., Jamil, A., Bai, Y., Ho, N., Zhao, M., Liu, Y., O’Connell, M.P., et al. (2016). Elevated basal serum tryptase identifies a multisystem disorder associated with increased TPSAB1 copy number. Nat Genet 48, 1564–1569.

MacArthur, J., Bowler, E., Cerezo, M., Gil, L., Hall, P., Hastings, E., Junkins, H., McMahon, A., Milano, A., Morales, J., et al. (2017). The new NHGRI-EBI Catalog of published genome-wide association studies (GWAS Catalog). Nucleic Acids Res 45, D896–D901.

Masamha, C.P., Xia, Z., Yang, J., Albrecht, T.R., Li, M., Shyu, A.B., Li, W., and Wagner, E.J. (2014). CFIm25 links alternative polyadenylation to glioblastoma tumour suppression. Nature 510, 412– 416.

Matoulkova, E., Michalova, E., Vojtesek, B., and Hrstka, R. (2012). The role of the 3’ untranslated region in post-transcriptional regulation of protein expression in mammalian cells. RNA Biol 9, 563–576.

Mayr, C. (2017). Regulation by 3’-Untranslated Regions. Annu Rev Genet 51, 171-194.

Mayr, C. (2018). What Are 3’ UTRs Doing? Cold Spring Harb Perspect Biol. pii: a034728

Muller-McNicoll, M., Botti, V., de Jesus Domingues, A.M., Brandl, H., Schwich, O.D., Steiner, M.C., Curk, T., Poser, I., Zarnack, K., and Neugebauer, K.M. (2016). SR proteins are NXF1 adaptors that link alternative RNA processing to mRNA export. Genes Dev 30, 553–566.

Musunuru, K., Strong, A., Frank-Kamenetsky, M., Lee, N.E., Ahfeldt, T., Sachs, K.V., Li, X., Li, H., Kuperwasser, N., Ruda, V.M., et al. (2010). From noncoding variant to phenotype via SORT1 at the 1p13 cholesterol locus. Nature 466, 714–719.

Park, H.J., Ji, P., Kim, S., Xia, Z., Rodriguez, B., Li, L., Su, J., Chen, K., Masamha, C.P., Baillat, D., et al. (2018). 3’ UTR shortening represses tumor-suppressor genes in trans by disrupting ceRNA crosstalk. Nat Genet 50, 783–789.

Pickrell, J.K. (2014). Joint analysis of functional genomic data and genome-wide association studies of 18 human traits. Am J Hum Genet 94, 559–573.

Shabalin, A.A. (2012). Matrix eQTL: ultra fast eQTL analysis via large matrix operations. Bioinformatics 28, 1353–1358.

Sheng, G., dos Reis, M., and Stern, C.D. (2003). Churchill, a zinc finger transcriptional activator, regulates the transition between gastrulation and neurulation. Cell 115, 603–613.

Stacey, S.N., Sulem, P., Jonasdottir, A., Masson, G., Gudmundsson, J., Gudbjartsson, D.F., Magnusson, O.T., Gudjonsson, S.A., Sigurgeirsson, B., Thorisdottir, K., et al. (2011). A germline variant in the TP53 polyadenylation signal confers cancer susceptibility. Nature Genetics 43, 1098.

Stegle, O., Parts, L., Piipari, M., Winn, J., and Durbin, R. (2012). Using probabilistic estimation of expression residuals (PEER) to obtain increased power and interpretability of gene expression analyses. Nature Protocols 7, 500.

Thomas, L.F., and Sætrom, P. (2012). Single nucleotide polymorphisms can create alternative polyadenylation signals and affect gene expression through loss of microRNA-regulation. PloS Comput Biol 8, e1002621.

Tian, B., and Manley, J.L. (2016). Alternative polyadenylation of mRNA precursors. Nature reviews Molecular cell biology. 18, 18–30

Urbut, S.M., Wang, G., Carbonetto, P., and Stephens, M. (2018). Flexible statistical methods for estimating and testing effects in genomic studies with multiple conditions. Nat Genet.

van der Maarel, S.M., Tawil, R., and Tapscott, S.J. (2011). Facioscapulohumeral muscular dystrophy and DUX4: breaking the silence. Trends in Molecular Medicine 17, 252–258.

Wang, S., Adrianto, I., Wiley, G.B., Lessard, C.J., Kelly, J.A., Adler, A.J., Glenn, S.B., Williams, A.H., Ziegler, J.T., Comeau, M.E., et al. (2012). A functional haplotype of UBE2L3 confers risk for systemic lupus erythematosus. Genes & Immunity 13, 380–387.

Weng, L., Li, Y., Xie, X., and Shi, Y. (2016). Poly(A) code analyses reveal key determinants for tissue-specific mRNA alternative polyadenylation. RNA 22, 813–821.

Xia, Z., Donehower, L.A., Cooper, T.A., Neilson, J.R., Wheeler, D.A., Wagner, E.J., and Li, W. (2014). Dynamic analyses of alternative polyadenylation from RNA-seq reveal a 3’-UTR landscape across seven tumour types. Nat Commun 5, 5274.

Yang, R., Gaidamakov, S.A., Xie, J., Lee, J., Martino, L., Kozlov, G., Crawford, A.K., Russo, A.N., Conte, M.R., Gehring, K., et al. (2011). La-related protein 4 binds poly(A), interacts with the poly(A)-binding protein MLLE domain via a variant PAM2w motif, and can promote mRNA stability. Mol Cell Biol 31, 542–556.

Yoon, O.K., Hsu, T.Y., Im, J.H., and Brem, R.B. (2012). Genetics and regulatory impact of alternative polyadenylation in human B-lymphoblastoid cells. PloS Genet 8, e1002882.

Zhang, H., Lee, J.Y., and Tian, B. (2005). Biased alternative polyadenylation in human tissues. Genome biology 6, R100.

Zhernakova, D.V., de Klerk, E., Westra, H.J., Mastrokolias, A., Amini, S., Ariyurek, Y., Jansen, R., Penninx, B.W., Hottenga, J.J., Willemsen, G., et al. (2013). DeepSAGE reveals genetic variants associated with alternative polyadenylation and expression of coding and non-coding transcripts. PloS Genet 9, e1003594.

## References

Bulik-Sullivan, B.K., Loh, P.R., Finucane, H.K., Ripke, S., Yang, J., Schizophrenia Working Group of the Psychiatric Genomics, C., Patterson, N., Daly, M.J., Price, A.L., and Neale, B.M. (2015). LD Score regression distinguishes confounding from polygenicity in genome-wide association studies. Nat Genet 47, 291–295.

Dobin, A., Davis, C.A., Schlesinger, F., Drenkow, J., Zaleski, C., Jha, S., Batut, P., Chaisson, M., and Gingeras, T.R. (2013). STAR: ultrafast universal RNA-seq aligner. Bioinformatics 29, 15–21.

Durinck, S., Spellman, P.T., Birney, E., and Huber, W. (2009). Mapping identifiers for the integration of genomic datasets with the R/Bioconductor package biomaRt. Nat Protoc 4, 1184– 1191.

Lindenbaum, P., and Redon, R. (2018). bioalcidae, samjs and vcffilterjs: object-oriented formatters and filters for bioinformatics files. Bioinformatics 34, 1224–1225.

Pruim, R.J., Welch, R.P., Sanna, S., Teslovich, T.M., Chines, P.S., Gliedt, T.P., Boehnke, M., Abecasis, G.R., and Willer, C.J. (2010). LocusZoom: regional visualization of genome-wide association scan results. Bioinformatics 26, 2336–2337.

Purcell, S., Neale, B., Todd-Brown, K., Thomas, L., Ferreira, M.A., Bender, D., Maller, J., Sklar, P., de Bakker, P.I., Daly, M.J., et al. (2007). PLINK: a tool set for whole-genome association and population-based linkage analyses. Am J Hum Genet 81, 559–575.

Quinlan, A.R., and Hall, I.M. (2010). BEDTools: a flexible suite of utilities for comparing genomic features. Bioinformatics 26, 841–842.

Stegle, O., Parts, L., Piipari, M., Winn, J., and Durbin, R. (2012). Using probabilistic estimation of expression residuals (PEER) to obtain increased power and interpretability of gene expression analyses. Nat Protoc 7, 500–507.

Storey, J.D., and Tibshirani, R. (2003). Statistical significance for genomewide studies. Proc Natl Acad Sci U S A 100, 9440–9445.

Urbut, S.M., Wang, G., Carbonetto, P., and Stephens, M. (2019). Flexible statistical methods for estimating and testing effects in genomic studies with multiple conditions. Nat Genet 51, 187– 195.

